# The RNA chaperone protein CspA stimulates translation during cold acclimation by promoting the progression of the ribosomes

**DOI:** 10.1101/2021.05.24.445485

**Authors:** Anna Maria Giuliodori, Riccardo Belardinelli, Melodie Duval, Raffaella Garofalo, Emma Schenckbecher, Vasili Hauryliuk, Eric Ennifar, Stefano Marzi

## Abstract

CspA is an RNA binding protein expressed during cold-shock in *Escherichia coli,* capable of stimulating translation of several mRNAs – including its own – at low temperature. We used reconstituted translation systems to monitor the effects of CspA on the different steps of the translation process and probing experiments to analyze the interactions with its target mRNAs. We specifically focused on *cspA* mRNA which adopts a cold-induced secondary structure at temperatures below 20°C and a more closed conformation at 37°C. We show that at low temperature CspA specifically promotes the translation of the mRNA folded in the conformation less accessible to the ribosome (37°C form). CspA interacts with its mRNA without inducing large structural rearrangement, does not bind the ribosomal subunits and is not able to stimulate the formation of the translation initiation complexes. On the other hand, CspA promotes the progression of the ribosomes during translation of its mRNA at low temperature and this stimulation is mRNA structure-dependent. A similar structure-dependent mechanism may be responsible for the CspA- dependent translation stimulation observed with other probed mRNAs, for which the transition to the elongation phase is progressively facilitated during cold acclimation with the accumulation of CspA.

## INTRODUCTION

Cold is a physical stress that influences conformation, flexibility, topology and interactions of every macromolecule in the cell. When subjected to abrupt temperature downshifts, mesophilic bacteria stop growing for several minutes until acclimation is established and growth resumes at lower temperature (for a review, see Giuliodori, 2016; Barria et al., 2013; Gualerzi et al., 2003; Weber and Marahiel, 2003). This cold acclimation phase is accompanied by drastic reprogramming of gene expression: whereas RNA, protein and lipid synthesis rates are in general reduced, the production of a small set of cold-shock (CS) proteins transiently increases (Giuliodori, 2016; Gualerzi et al., 2003; Phadtare and Inouye, 2004). These CS proteins are mainly transcription and translation factors as well as proteins involved in RNA structure remodeling, such as RNA chaperones, RNA helicases, and exoribonucleases (Jones et al., 1996; Bae et al., 2000; Yamanaka and Inouye, 2001; Gualerzi et al., 2003; Cairrão et al., 2003). The induction of CS gene expression is done at both the transcriptional and post-transcriptional levels (Gualerzi et al., 2003). Surprisingly, an increase in mRNA abundance after cold-shock does not necessarily lead to an increased synthesis of the corresponding protein (Goldenberg et al., 1997). The likely explanation is that low temperature impairs translation, affecting both the initiation (Broeze et al. 1978; Farewell and Neidhardt, 1998; Zhang et al., 2018) and the elongation (Friedman and Weinstein, 1964; Farewell and Neidhardt, 1998; Zhang et al., 2018) phases. The ability of the translational machinery to synthesize proteins under these unfavorable conditions relies on *cis*-acting elements encoded in mRNAs, whose existence was demonstrated in cells (Mitta et al., 1997; Yamanaka et al., 1999; Etchegaray and Inouye, 1999; Zhang et al., 2018) and using *in vitro* assays (Giuliodori et al., 2010, Giuliodori et al, 2019). *Trans*-acting factors play an important role in this process, such as the initiation factors IF1 and IF3, and the protein CspA, whose levels specifically increase during cold-shock (Giuliodori et al., 2004; Giuliodori et al., 2007; Di Pietro et al., 2013; Zhang et al., 2018). The timing of cold-shock gene induction suggests that the expression of some CS genes might be dependent on the synthesis of early CS proteins (Weber and Marahiel, 2003; Zhang et al., 2018).

CspA, a member of the CS protein (Csp) family, is the most well-studied *E. coli* CS protein (Yamanaka et al., 1998). Out of the nine paralogues, seven are cold-inducible (CspA, CspB, CspE, CspF, CspG, CspH and CspI) and two are expressed only at 37°C (CspC and CspD) (Giuliodori 2016, Zhang et al, 2018). Furthermore, expression of CspF and CspH is also induced upon urea challenge (Withman et al., 2013). To generate a cold-sensitive phenotype, four out of the nine *csp* genes must be knocked out in *E. coli* genome, and this cold-sensitive phenotype can be reverted by overexpressing any of the *csp* members, with the exception of *cspD* (Xia et al., 2001). CspA is a small protein of 70 amino acids comprised of a single OB fold domain, similar to S1 domain (Newkirk et al., 1994; Schindelin et al., 1994). The protein preferentially binds single strand regions of RNA and DNA (Jiang et al, 1997), but the sequence specificity of the interaction is still unclear. While some reports show preferential binding of CspA to polypyrimidine-rich sequences (Lopez and Makhatadze, 2000), others studies suggest that CspA lacks sequence specificity (Jiang et al., 1997). Importantly, the 5′-UTR *cspA* mRNA encoding CspA was suggested to be a *bona fide* target of CspA-mediated regulation (Jiang et al. 1997). In *E. coli* the extent of cold shock induction of *cspA* mRNA is growth phase-dependent (Brandi et al., 1999). When cells are subjected to cold-shock during mid-late exponential growth, they abundantly transcribe and translate *cspA* mRNA *de novo* to increase the level of the protein. Conversely, when cells are subjected to cold-shock in the early stage of growth, they use CspA and its transcript synthetized at 37°C predominantly, at least at the beginning of the cold adaptation phase.

While several structures of Csp proteins are available – namely, *E. coli* CspA (Newkirk et al., 1994; Schindelin et al., 1994), *B. subtilis* CspB (Schindelin et al., 1993; Schnuchel et al., 1993), *B. caldolyticus* CspB (Mueller et al., 2000), and *Thermotoga maritima* CspB (Kremer et al., 2001) – only two structures of RNA-Csp protein complexes have been determined. Specifically, the crystal structure of *B. subtilis* CspB – the major cold shock protein in this bacterium *–* has been solved in complex with two oligoribonucleotides: 5′-UUUUUU-3′ and 5′-GUCUUUA-3′ providing a framework to explain how the OB fold domain interacts with RNA (Sachs et al., 2012). Seven aromatic residues and two lysines located on the surface of CspB are the key players in binding to RNA.

Even if the rules governing the CspA-RNA interaction are not known, its low binding selectivity and affinity (association constant in the μM range (Jiang et al., 1997; Phadtare and Inouye, 1999; Lopez and Makhatadze, 2000) are typical of RNA chaperone proteins, which bind RNAs only transiently (Mayer et al., 2007; Rajkowitsch and Schroeder, 2007; Duval et al., 2017). However, CspA is a highly abundant protein during cold stress: it accounts for up to 10% of the total proteins during cold adaptation (Brandi et al., 1999), with intracellular CspA concentration reaching 100 μM (Bae et al., 1999; Brandi et al., 1999). Therefore, it is expected that in cold-stressed cells several molecules of CspA could bind simultaneously to an individual target mRNA (Ermolenko and Makhatadze, 2002; Zhang et al., 2018). Given CspA propensity to bind single strand (ss) regions in nucleic acids, the protein is expected to play an important role in modulating mRNA structures induced/stabilized by low temperatures, thus playing crucial role in regulating their expression. This hypothesis is supported by experiments showing that (i) CspA promotes melting of the secondary structure of MS2 mRNA (Phadtare et al., 2009), (ii) CspA acts as both transcriptional (La Teana et al., 1991; Jones et al., 1992) and translational (Giuliodori et al., 2004) activator in the cold, and (iii) CspA acts as an RNA chaperone (Jiang et al., 1997; Rennella et al., 2017; Zhang et al., 2018).

Several crucial questions regarding the cellular functions of CspA and its mechanism of action still remain to be answered. As, for example, the determinants for CspA activation of specific target genes and the mechanism for CspA-dependent stimulation of their expression. In the present work, we have characterized the mechanism of translational activation of *cspA* mRNA by CspA using a cell-free reconstituted translation system and probing methods. The translation efficiency of the two forms of *cspA* mRNA was compared: the newly synthesized mRNA adopts an open, cold-induced, conformation below 20°C and a more closed conformation at 37°C, which is stabilized at low temperatures (Giuliodori et al., 2010). Using the two *cspA* mRNA forms in translation assays performed at low temperature, we demonstrate that CspA specifically promotes translation of the mRNA with the more closed structure. Combining crosslinking and footprinting approaches, we demonstrate that CspA binds to the two *cspA* mRNA conformations at various positions. However, CspA binding is neither able to unwind the structures of the stably folded mRNA nor capable of promoting the formation of translation initiation complexes. Our experiments suggest that at low temperature, CspA assists the progression of ribosomes along its highly structured mRNA. Furthermore, we demonstrate by cross-linking experiments that CspA binds to other mRNAs preferentially in a position located downstream from the initiation codon and can stimulate the translation of some of the bound transcripts. Indeed, analysis of available ribosome profiling data during cold acclimation shows that these CspA-dependent mRNAs present ribosomes stalled on the initiation codons, which progress to translation elongation with the cellular accumulation of CspA. Eventually, we proposed a model that takes into account these results and explain the translation activity of CspA upon cold shock.

## RESULTS

### CspA favors the translation of less favorable structure of cspA mRNA during cold shock

To uncover the possible effects of cold-shock trans-acting factors on the translation of mRNAs with unfavorable secondary structures at low temperature, we first used a translation cell-free system using crude S30 extracts (i.e. a bacterial content deprived from membrane debris) prepared from cells grown at 37°C (control) or exposed to 15°C for 120 minutes (cs extracts). The latter type of extract contains high levels of cold-shock proteins synthesized when cells reprogram their genetic expression after sensing the cold (Giuliodori, 2016). *In vitro* translation reactions were programmed with *cspA* mRNA, which acts as useful tool for studying structural transitions in RNA and the role of CS factors. After denaturation at 90°C, this transcript can be refolded in two different structures: the cold-structure, which exists only at a temperature below 20°C, and the 37°C-structure (Giuliodori et al., 2010). While the cold-structure is competent in efficient recruitment of 30S ribosomal subunits to AUG initiation codon (Figure 1A), in the 37°C-form both the Shine-Dalgarno sequence (SD) and the initiation codon are partially occluded (Figure 1B). Notably, the 37°C-form is very stable and is maintained and stabilized upon incubation at low temperature (Giuliodori et al., 2010).

**FIGURE 1.**
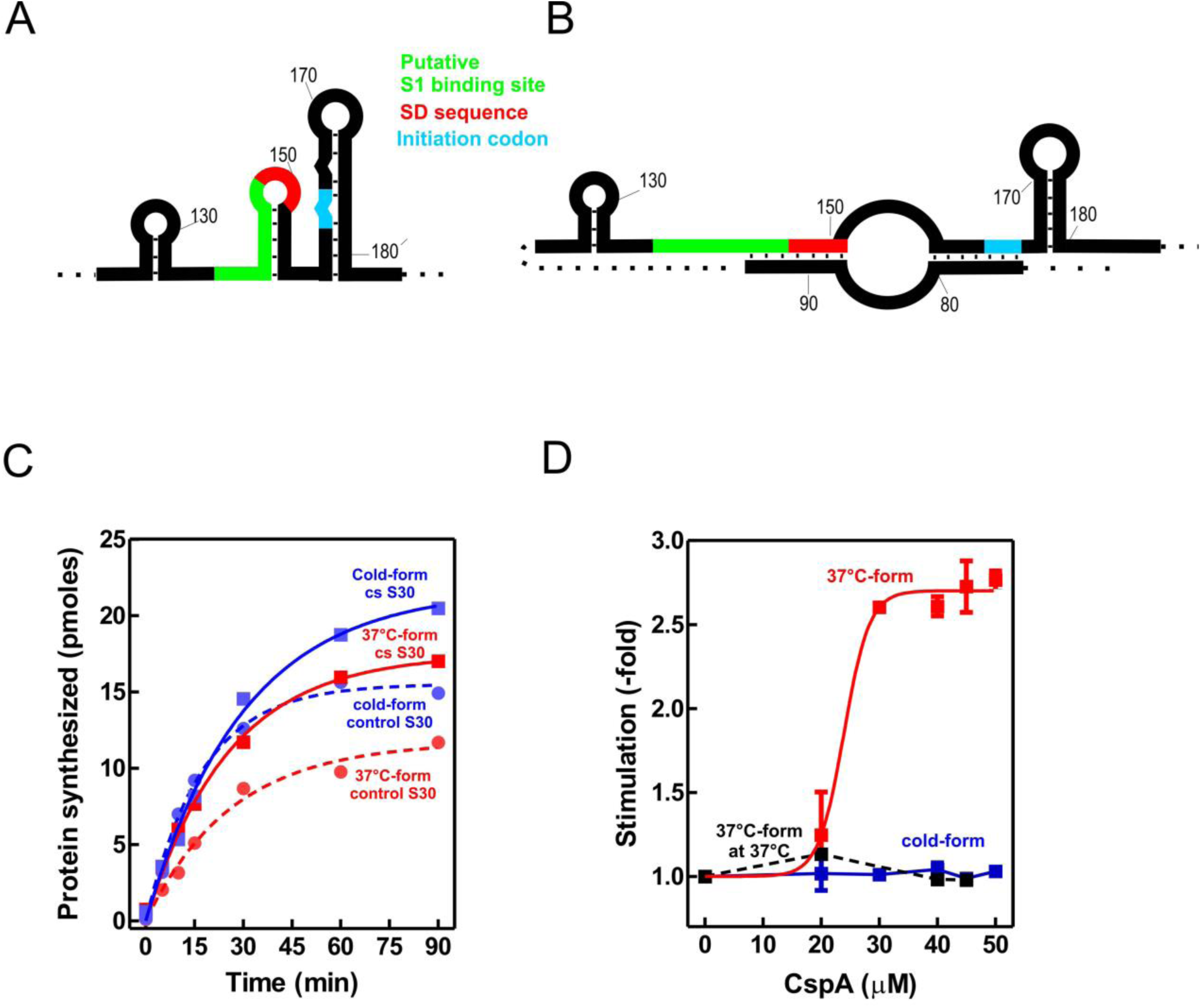
Effect of CspA on *cspA* mRNA translation at low temperature. Schematic representation of the secondary structures of the Translation Initiation Region (TIR) of the A) cold-form and B) 37°C of *cspA* mRNA. The SD sequence, the start codon and the putative S1-binding site are indicated in red, light blue and green, respectively (Giuliodori et al., 2010). (C) *in vitro* translation at 15°C with control (dashed lines) and cold-shock (solid lines) S30 extracts; the experiments were carried out with *cspA* mRNA folded in the cold-form (blue symbols) or in the 37°C-form (red symbols). (D) *in vitro* translation with control 70S and S100 in the presence of the indicated amounts of purified CspA at 15°C (solid line) with the cold-form (blue) or the 37°C-form (red) of *cspA* mRNA. The reaction was also performed at 37°C only with the 37°C-form (dashed black line). Data points in panel B are the average of two independent experiments. Error bars represent the standard deviations. Further details are given in Materials and Methods.

The translational activities of the two *cspA* mRNA forms were tested with the two types of cell extracts at 15°C (Fig. 1C). The data show that the cold structure is more efficiently translated than the 37°C-structure. However, during the first 30 minutes of the time-course, the extracts from cold-shocked cells translate with similar efficiency both forms of *cspA* mRNA. This suggests that the intracellular milieu of cold-treated *E. coli* is enriched in factors that support the translation of the mRNA with the less favorable secondary structure. Since the most abundant protein in the cold-shock extract – CspA – is able to stimulate protein synthesis at low temperature (Giuliodori et al., 2004), we next investigated its role in translation. To this end, translation of the cold- and 37°C-forms of *cspA* mRNA was studied in the presence of increasing amounts of purified CspA using 70S ribosomes and post-ribosomal supernatant (S100) prepared from cells that were not exposed to low temperature (Fig. 1D). The reactions were carried out at 15°C with the two forms of *cspA,* whereas at 37°C only the activity of the 37°C-structure of *cspA* mRNA could be probed, as the cold structure exists only at temperatures below 20°C (Giuliodori et al., 2010). Our results (Fig. 1D) clearly demonstrate that CspA strongly promotes (> 2.5- fold) the translation of the less-favorable 37°C-structure of *cspA* mRNA at low temperature, while it does not affect the translation of the other and more open *cspA* mRNA form. The effect is strongly dose-dependent, as the translation sharply increased when CspA concentration rises above 20-25 µM. Interestingly, this stimulatory activity of CspA is not observed at 37°C.

### CspA assists the progression of the ribosome along the structured mRNA at low temperature

To investigate the mechanism by which CspA stimulates the translation process, we tested the effect of purified CspA on the recruitment of the mRNA conformers to the 30S subunit either in the presence (Fig. 2A) or in the absence (Fig. 2B) of IFs and the initiator fMet-tRNA_i_. We established that CspA does not assist the binding of its mRNA to the small ribosome subunit, and we confirmed that the cold-form mRNA binds better (approximately 2.5-fold) to the 30S subunits than the 37°C-form mRNA (Giuliodori et al., 2010). Next, using filter binding (Fig. 2C) and toeprinting assays (Fig. 2D), we probed the ability of CspA to promote the binding of fMet-tRNA_i_^Met^ to the 30S subunits and the consequent formation of the active initiation complexes in the presence of the 37°C-form of *cspA* mRNA at low temperature. The two experiments showed that the assembly of the initiation complex is insensitive to the addition of CspA.

**FIGURE 2.**
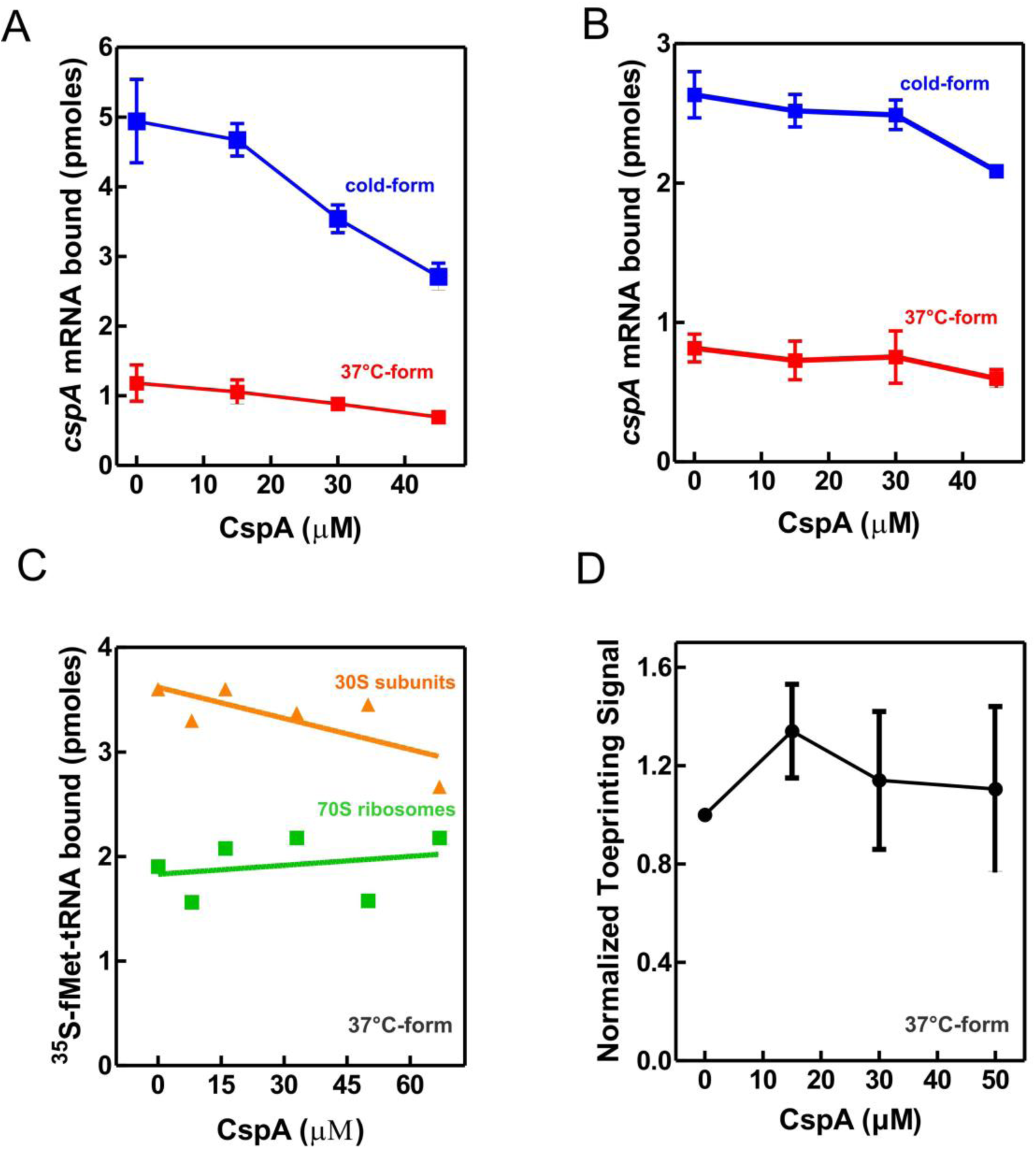
Effect of CspA on the individual steps of translation initiation. The effect of increasing amounts of CspA on the binding of *cspA* mRNA on the 30S subunits was monitored at 15°C by filter binding of ^32^P-labelled *cspA* mRNAs folded in the cold-form (blue) or in the 37°C-form (red) in the absence (A) or in the presence (B) of IFs and fMet-tRNA_i_^Met^. The effect of increasing amount of CspA on the initiation phase of translation at 15°C was also investigated by analyzing: (C) the binding of ^35^S-fMet-tRNA_i_^Met^ to 30S subunits (green squares) and to 70S ribosomes (orange triangles) programmed with IFs and the 37°C-form of *cspA* mRNA by filter binding assay and (D) the localization of 30S on the translation start site of *cspA* mRNA by toeprint assay. The data points in panels A, and B are the average of triplicates. The data points in panel D result from the quantification of toeprint signals of two independent experiments. Error bars represent the standard deviations. Further details are given in Materials and Methods.

We next explored whether this protein could stimulate the translation elongation step in the cold. To this end, we have developed a test system based on the *E. coli* RelE toxin, which cleaves between the second and the third nucleotide of the mRNA codon in the ribosomal A site in the absence of the cognate A-site tRNA (Pedersen et al., 2003; Neubauer et al., 2009), in a so-called “RelE walking” experiment. Radiolabeled cold- and 37°C-forms of *cspA* mRNA were translated *in vitro* at 15°C using the PURE system (NEB) – a reconstituted system of the *E. coli* translation machinery with reduced concentration of charged asparagine tRNA (Asn-tRNA^Asn^) (Shimizu at al., 2001 and patent US7118883b2). At the end of the incubation, chloramphenicol and RelE were added to the reaction mixtures to stabilize the polysomes and to cut the mRNA at the codons in which the ribosomes were blocked due to the low content of Asn-tRNA^Asn^, respectively. Using polyacrylamide gel electrophoresis (PAGE), we monitored the extent of RelE cleavages on the three Asn triplets AAC (13^th^, 39^th^ and 66^th^ codon) and on the first A-site codon after the AUG. This experiment provides new data concerning the fraction of ribosomes engaged in mRNA translation and the ribosomal progression along the transcript.

Figure 3A shows that RelE cleaves extensively all Asn codons of the cold *cspA* mRNA form, independently of CspA, confirming its translability at low temperature. Interestingly, the intensities of the RelE cleavages detected with the 37°C-form of *cspA* mRNA (Fig. 3B) are much weaker compared to those of the cold form, the only exception being the cuts at the first A-site codon, which are comparable in the two forms of *cspA* mRNA. The data suggest that the number of ribosomes transiting along *cspA* mRNA and pausing at the asparagine codons is significantly lower in the case of the highly structured 37°C-form than in the cold-form of the mRNA. Notably, the addition of CspA to the translation system programmed with the 37°C-form causes intensification of the RelE cleavages, suggesting that the number of elongating ribosomes has enhanced (Fig. 3B). Indeed, quantification of the gel bands (Fig. 3C and 3D, normalized values) reveals that CspA induces on average a 2.5-fold increase of progression with the 37°C-form, a value very close to the observed stimulatory effect on translation (Fig. 1D). The fact that CspA does not affect the rate of RelE cleavage of the cold-form of *cspA* mRNA (Fig. 3C) excludes the possibility that CspA could influence directly the RelE activity on the 37°C- form. The fact that the 3′ termini of the two *cspA* mRNAs are identical (Giuliodori et al., 2010) excludes the possibility that the diverse RelE cleavage rates between the two forms could be promoted by a different rate of ribosome recycling. Finally, the RelE cuts at the A- site of the 70S initiation complex are similar in the absence and in the presence of CspA, thus ruling out the possibility that CspA could favor the occupancy of the A site by the aa- tRNA in the initiation phase.

**FIGURE 3.**
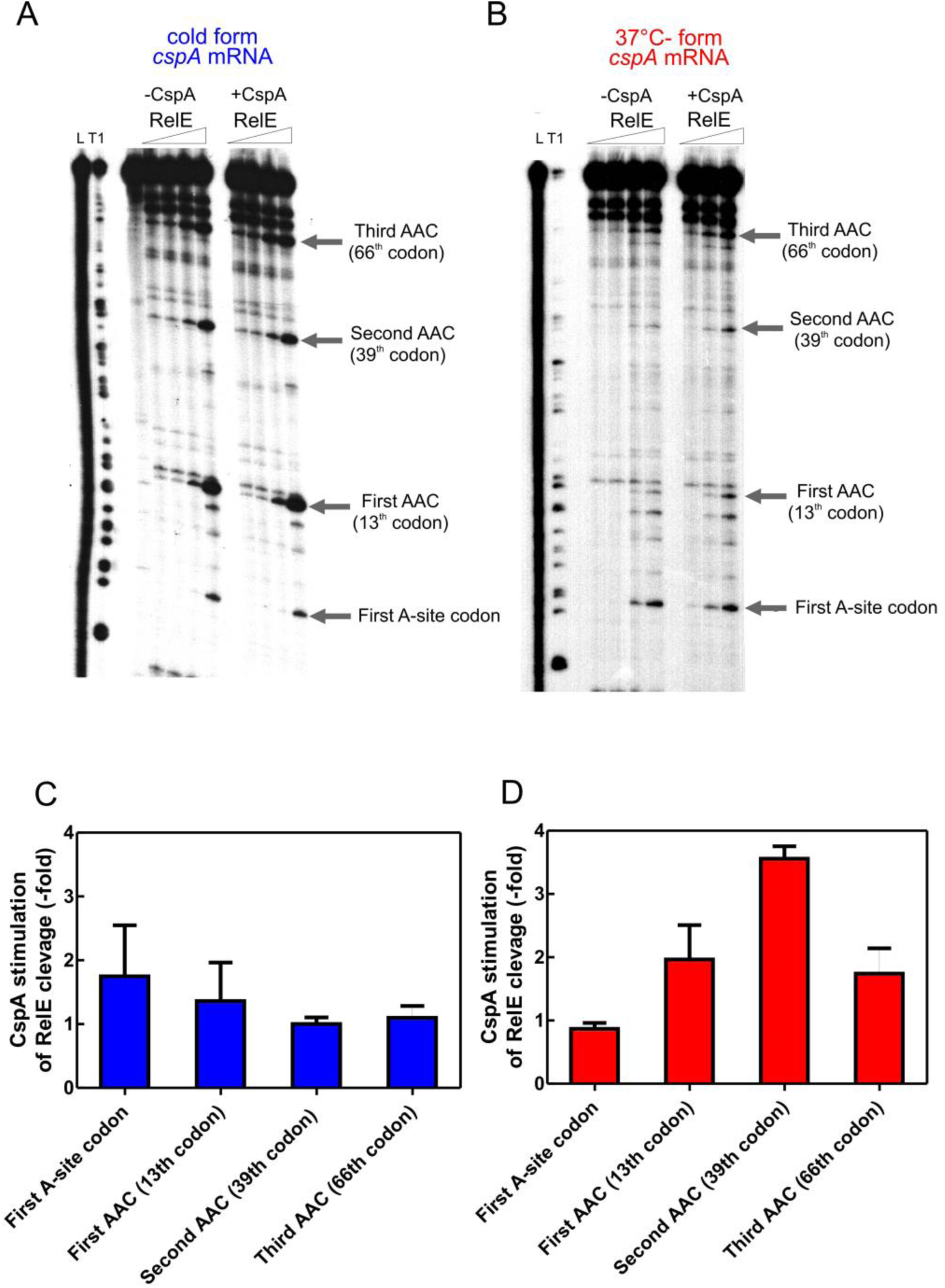
RelE walking experiment. ^32^P-labelled *cspA* mRNA, folded in the cold-structure (A) or in the 37°C-structure (B), was used as templates for an *in vitro* translation assay with the PURE system and then cleaved with 0, 0.16, 0.72 and 1.44 μM of RelE. Lane L: alkaline ladder; lane T1: RNase T1 ladder. The first A-site codon (after the AUG initiation codon) and the AAC codons specifying Asn are indicated. Numbering is given according to the initiation codon. C and D show the effect of CspA (30 µM) on the intensity of RelE cleavages (normalized to the total radioactivity present in each lane) on the 15°C- and 37°C-structures, respectively. Averages of the fold changes observed using the three different RelE concentrations are reported with standard deviations.

Based on the RelE walking experiments, we propose that CspA promotes progression of the ribosomes on structured mRNAs during translation elongation at low temperature.

### Binding of CspA to cspA mRNA is responsible for translation stimulation

Isothermal Titration Calorimetry (ITC) is a powerful technique for studying interactions between native proteins and their RNA targets (Klebe et al., 2015). We used this approach to probe the possible interaction of CspA with the 70S ribosome (Fig. 4A), individual 30S (Fig. 4B), and 50S subunits (Fig. 4C) at 15°C, 25°C, and 35°C (Fig. 4-figure supplement 1). We fail to detect a specific interaction between CspA with either the 70S ribosome or the isolated subunits. The small variations observed are due to the heat released by the disassembly of CspA or ribosome aggregates in buffer upon dilution (Fig. 4-figure supplement 1). On the other hand, as expected, we detected the specific binding of initiation factor IF1 to the 30S subunit at low temperature under comparable conditions (Kd = 806 nM). Notably, IF1 and CspA share impressive structural similarity (Gualerzi et al., 2011) and are both RNA binding proteins (Phadtare and Severinov, 2009); however, IF1 overexpression in *E. coli* does not suppress the defects of the *csp* quadruple deletion strain (Phadtare and Severinov, 2009).

**FIGURE 4.**
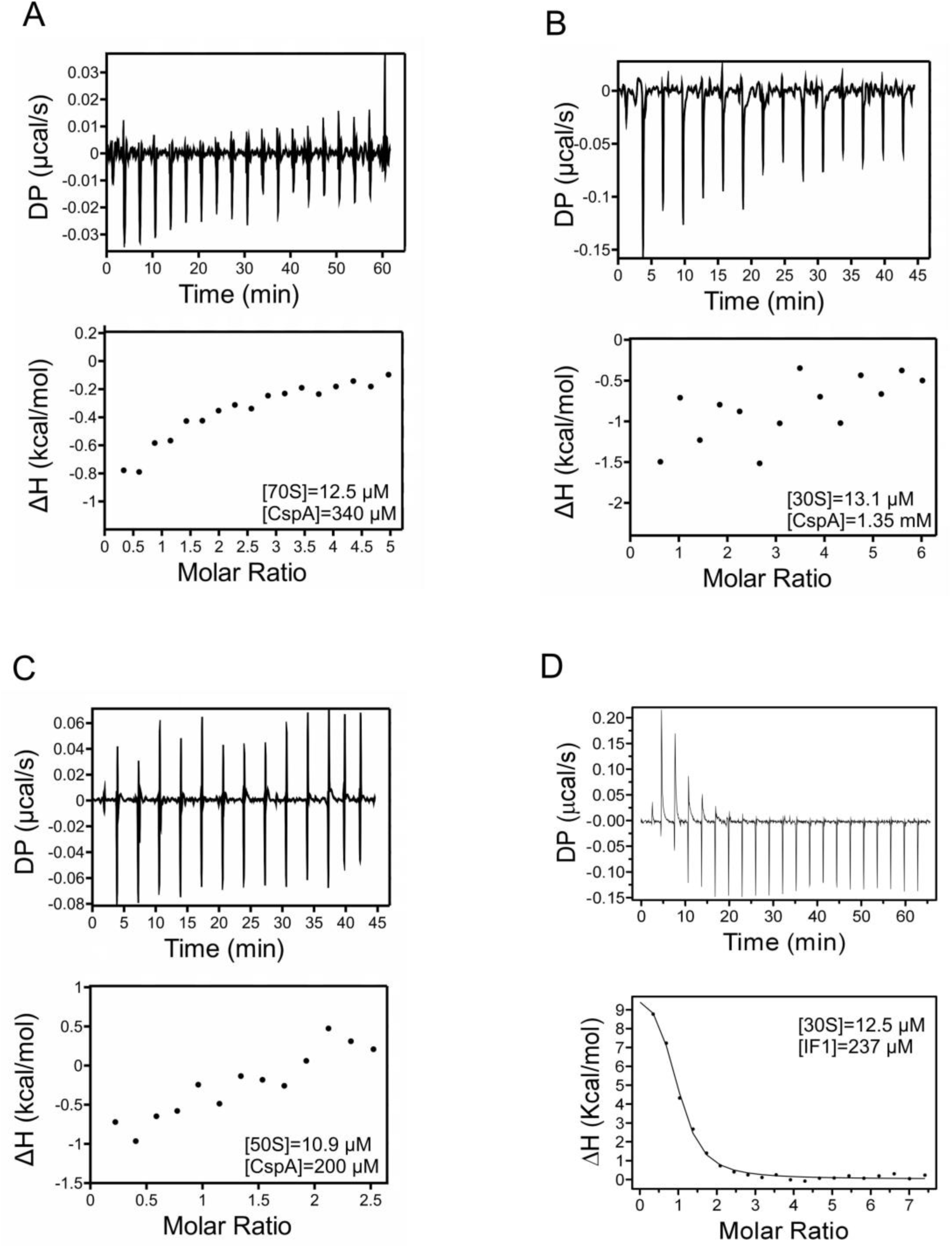
ITC analysis of CspA interaction with the 70S ribosomes and ribosomal subunits. Titration of *E. coli* ribosomes with CspA was studied at 15°C by sequential 2 µL injections of CspA in a cell containing 200 µL of 70S ribosomes (A), 30S subunits (B) or 50S subunits (C). Control experiments were performed using IF1 and 30S subunits at the indicated concentrations (D).

CspA has been shown to bind RNA including the 5′-UTR of its own transcript (Jiang et al. 1997). Therefore, we investigated CspA-*cspA* mRNA interaction using Electrophoresis Mobility Shift Essay (EMSA). Given the large size (428 nts) of the full-length *cspA* transcript and the small mass of CspA (7.4 KDa), separation of such complexes constituted a technical challenge. Therefore, the analysis was done using a *cspA* mRNA fragment of 187 nts (*187cspA* RNA) consisting of the whole 5′ UTR plus 27 nts of the coding region. Importantly, this fragment adopts a secondary structure highly similar to that found in the full-length transcript at low temperature (Giuliodori et al., 2010). Complex formation with increasing concentrations of CspA is shown in in Fig. 5A and B. Below 80 µM of CspA, a complex with minor gel retardation is observed (indicated with a thin arrow). However, as the amount of CspA exceeds 80 µM, a super-shifted band appears (indicated with a thick arrow), whose mobility continues to decrease with increasing amounts of CspA. This result indicates that the *187cspA* RNA contains multiple binding sites for CspA that are progressively occupied as the concentration of the protein rises. The appearance of the super-shift supports the hypothesis of a cooperative binding to RNA by CspA (Jiang et al., 1997; Lopez and Makhatadze, 2000). These experiments demonstrate that multiple CspA bind to its mRNA at both low (Fig. 5A) and high temperatures (Fig. 5B) and confirm that this protein can bind also structured RNA molecules, although in this case the multi-protein complexes are formed only at high protein concentrations.

**FIGURE 5.**
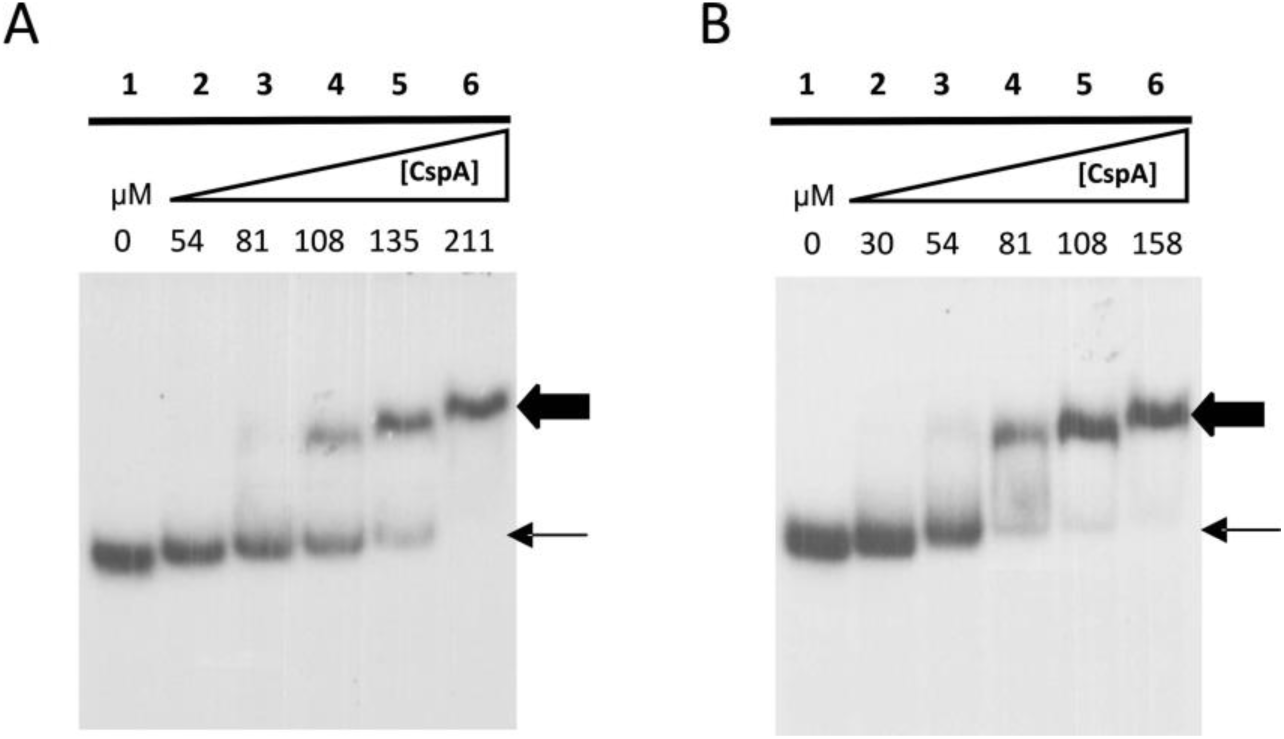
CspA-*187cspA* RNA interaction studied by EMSA. The binding of CspA to ^32^P-labeled *187cspA* RNA performed at 20°C (A) or at 37°C (B) was analyzed in gel retardation assays as described in Materials and Methods using the concentrations of protein indicated at the top of the gels. The thin and thick arrows indicate the complexes with high and low mobility, respectively.

The details of the CspA:*cspA* mRNA interaction at 15°C were then dissected using three different approaches: (i) UV-induced cross-linking and (ii) enzymatic probing, and Fe-EDTA footprinting. The resulting cross-link and footprint patterns are reported in the structure models of the cold-form (Fig. 6) and the 37°C-form (Fig. 7) of *cspA* mRNA, while the electrophoretic analyses are shown in Fig. 7-figures supplement 1, 2 and 3. In the cold-form, we identified 11 main sites, which were either protected or cross-linked to CspA. Overall, the CspA sites are mainly positioned in apical or internal loops, extending also into the adjacent helices. Most of these sites are rather large, especially sites 1, 7, 9 and 10, which are located at positions 12-36, 170-186, 266-281 and 321 to 337, respectively. Notably, the CspA-induced cross-links at sites 7, 9 and 10, which also overlap with CspA induced protections against enzymatic cleavages or FE-EDTA, are particularly strong.

**FIGURE 6.**
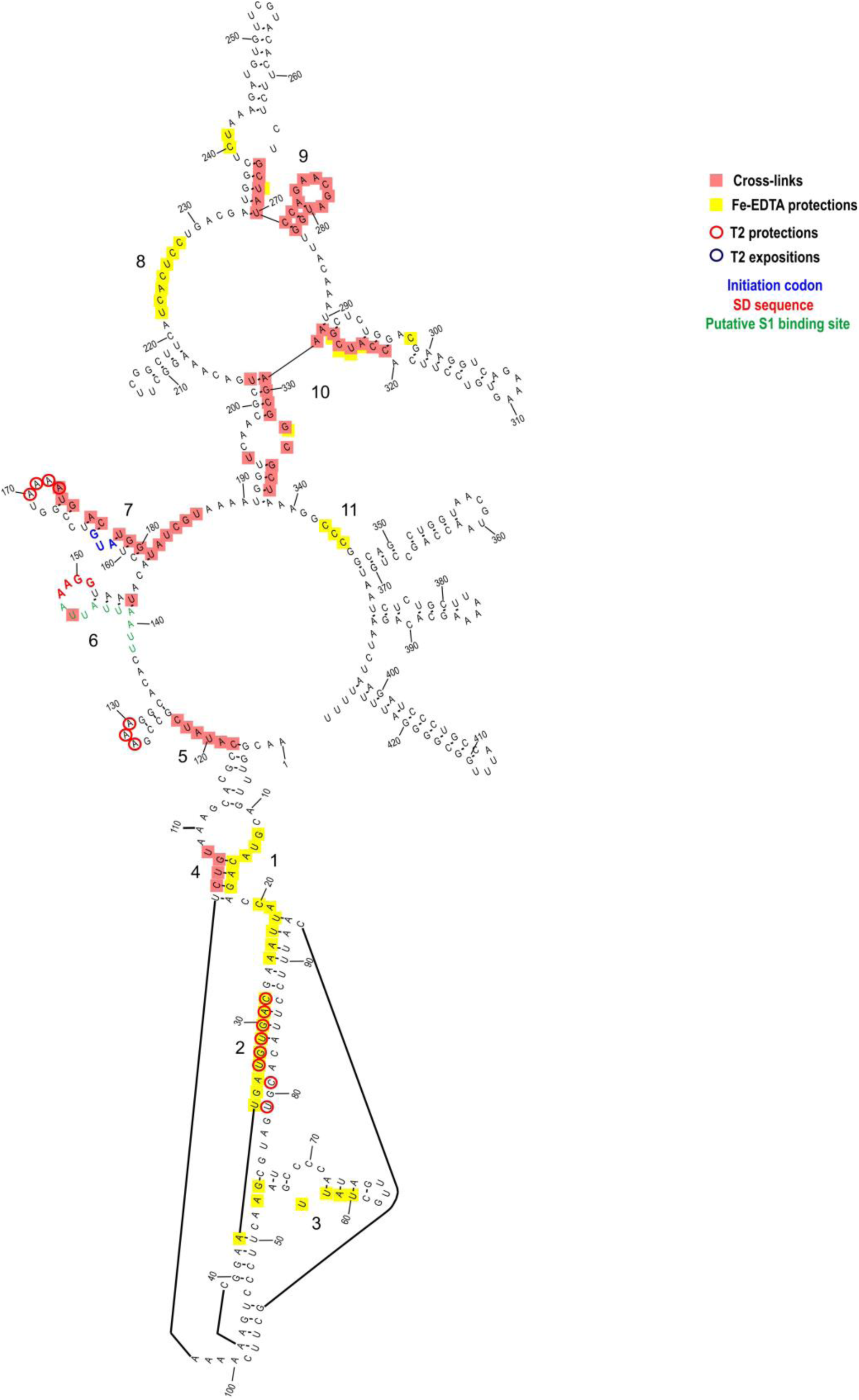
CspA-binding sites identified on the cold-form of *cspA* mRNA. The footprinted/crosslinked positions reported on the structural model derive from the probing experiments performed at 15°C using the *cspA* mRNA folded in the cold-conformation. The SD sequence (red), the start codon (blue) and the putative S1-binding site (green) are indicated. The secondary structure model of *cspA* mRNA is taken from Giuliodori et al., 2010.

**FIGURE 7.**
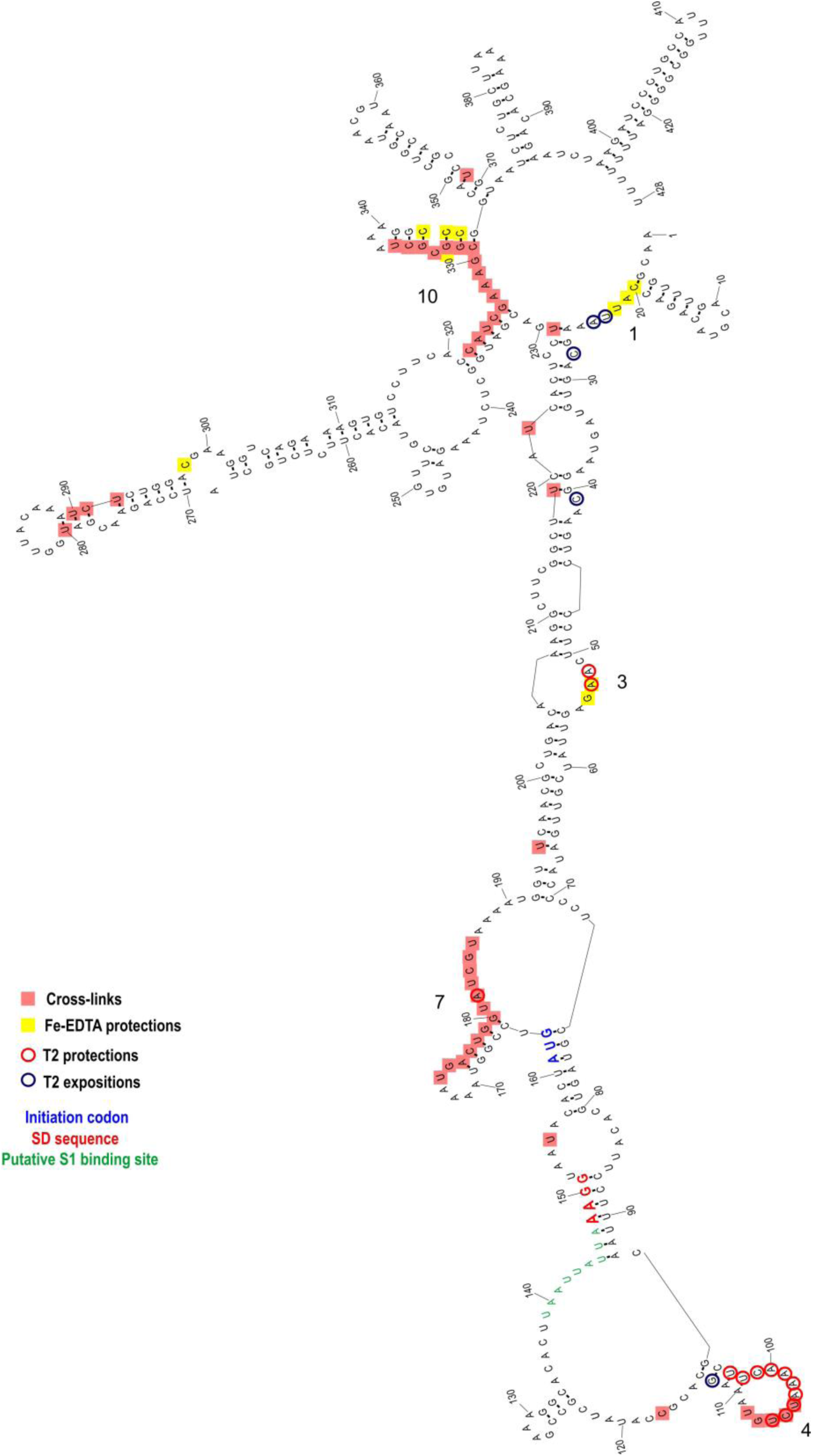
CspA binding sites identified on the 37°C-form of *cspA* mRNA. The footprinted/crosslinked positions reported on the structural model derive from the probing experiments performed at 15°C using the *cspA* mRNA folded in the 37°C-conformation. The SD sequence (red), the start codon (blue) and the putative S1-binding site (green) are indicated. The secondary structure model of *cspA* mRNA is taken from Giuliodori et al., 2010.

Probing the 37°C-form of *cspA* mRNA bound to CspA shows significantly different patterns as compared to the cold-form. Only 5 of the 11 sites present in the cold-form had a counterpart in the 37°C-form, namely sites 1, 3, 4, 7 and 10, while the other regions became insensitive to CspA (sites 2, 5, 6, 8, 9, and 11). Furthermore, the binding sites were shorter as compared to the cold-form, with the exception of site 7 at the beginning of the coding region, which remained quite extended. Finally, the cross-links were overall less intense and much more dependent upon CspA concentration than in the case of the cold-form.

The above-described differences can be likely attributed to the more compact structure of the 37°C-form, characterized by a long helix interrupted by several internal loops and bulged bases formed by the interaction between the 5′ UTR and part of the coding region (nucleotides C232 to G326). This closed conformation is further stabilized at low temperature (15°C) at which the probing experiments were performed. Indeed, the reduced binding of CspA to the 37°C-form mRNA is not surprising considering the preference of Csp proteins for single stranded nucleic acids. Most likely, CspA needs unstructured regions for the initial contacts with the target RNA.

### Binding of CspA to a short cspA mRNA fragment affects the mRNA conformation

The secondary structure of three *cspA* mRNA fragments of increasing length (i.e. 87, 137 and 187 nts) from the transcriptional start site was previously analysed (Giuliodori et al., 2010). These *cspA* mRNA fragments were designed as representative of RNA folding intermediates occurring during transcription. Their structures do not vary with temperature, and the 137*cspA* and 187*cspA* fragments adopt similar folding as in the full length cold-form of *cspA* mRNA. To investigate the role played by CspA on the initial mRNA folding process, we have analyzed the footprint of CspA on both the *87cspA* and the*137cspA* RNA fragments using RNase V1 (specific of double-stranded regions), RNase T1 (specific of unpaired guanine), and RNase T2 (specific of unpaired A>U>C). Binding of CspA to the short *87cspA* RNA significantly affects the RNase cleavage pattern, but only at a concentration of CspA above 50 µM (Fig. 8A). For instance, protections against RNase T2 were observed between A29 and U33, and at positions U58 and U76; concomitantly enhanced RNase cleavages were found at positions C41, C50, A51, G65 and C81-A82. On the other hand, the addition of high concentrations of CspA had only minor effects on the structure of *137cspA* RNA (Fig. 8B and supplemental 1). For instance, the CspA- dependent protections in the region A29 and U33 of *87cspA* RNA were no longer observed.

**FIGURE 8.**
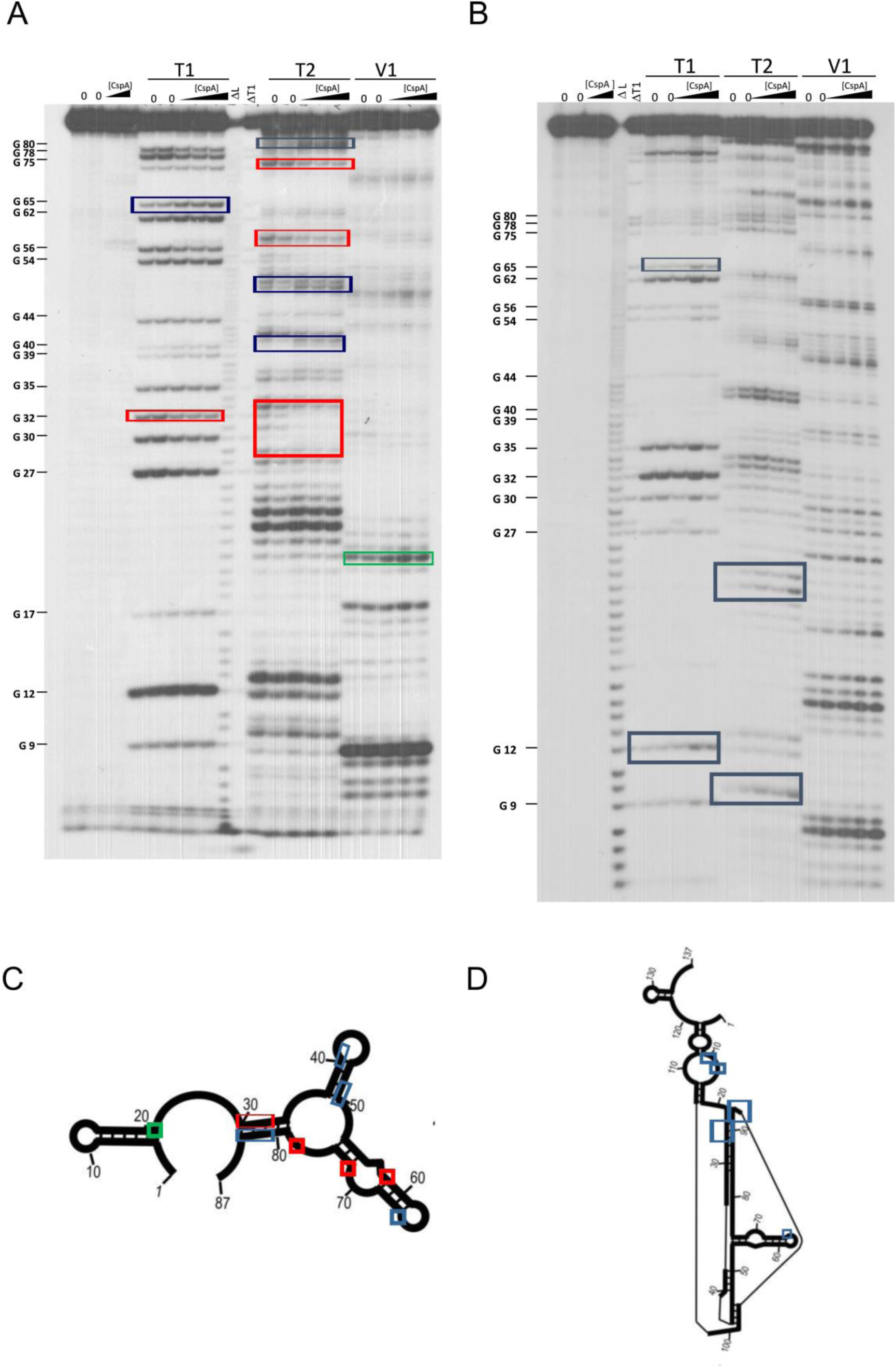
Footprinting experiments of fragments 87*cspA* and 137cspA RNAs. (A) Short electrophoretic migration of the fragments generated by RNase T1 (T1), RNase T2 (T2) or RNase V1 (V1) digestion of 5′-end [^32^P]-labelled (A) 87*cspA* RNA and (B) 137cspA RNA. The experiments were carried out in the absence (lanes 0) or in the presence of 51, 81, and 105 µM of CspA (increasing concentrations are indicated with a triangle). Lane ΔT: RNase T1 cleavages under denaturing conditions; lane ΔL: alkaline ladder. The red and blue boxes indicate the positions protected or exposed by CspA, respectively, while the green boxes indicate the V1 cuts enhanced by CspA. The same positions are reported on the schematic structural model (Giuliodori et al., 2010) of the 87cspA RNA (C) or of the 137cspA RNA (D).

These data suggest that CspA might have different functional impacts *in vivo* during the transcription process of cspA that will depend on many factors including CspA concentration, and kinetics of RNA transcription and folding.

### CspA preferentially binds to short RNA sequences containing YYR motif

Inspection of all sites covered by CspA revealed that 11 out of 15 of these regions comprise an YYR (pyrimidine-pyrimidine-purine) motif. Multiple alignments were performed using the YYR motif as the reference sequence (Fig. 9A). Although the motif is highly degenerated, the YYR motif seems not to be followed by a G two positions downstream from the R. The degree of specificity of CspA for the identified sequence features was then tested by tryptophan fluorescence titration experiments using an RNA oligonucleotide (Oligo1: 5′-AA**CUG**GUA-3′) whose sequence reflected the conserved positions as shown in Figure 9B. The experiment was also performed with 5 other RNA oligos in which each one of the bases located in the central positions of Oligo1 was individually replaced by A. In addition, a poly-A oligo was also used. The data (Fig. 9C and Table 1) show that the single nucleotide changes caused only small variations of the dissociation constant (K_D_ around 1 µM), with the exception of oligo 6 (G replaced by A at position 6), which produced a 5-fold increase of the K_D_. A similar decrease of affinity was observed with the poly-A oligonucleotide.

**FIGURE 9.**
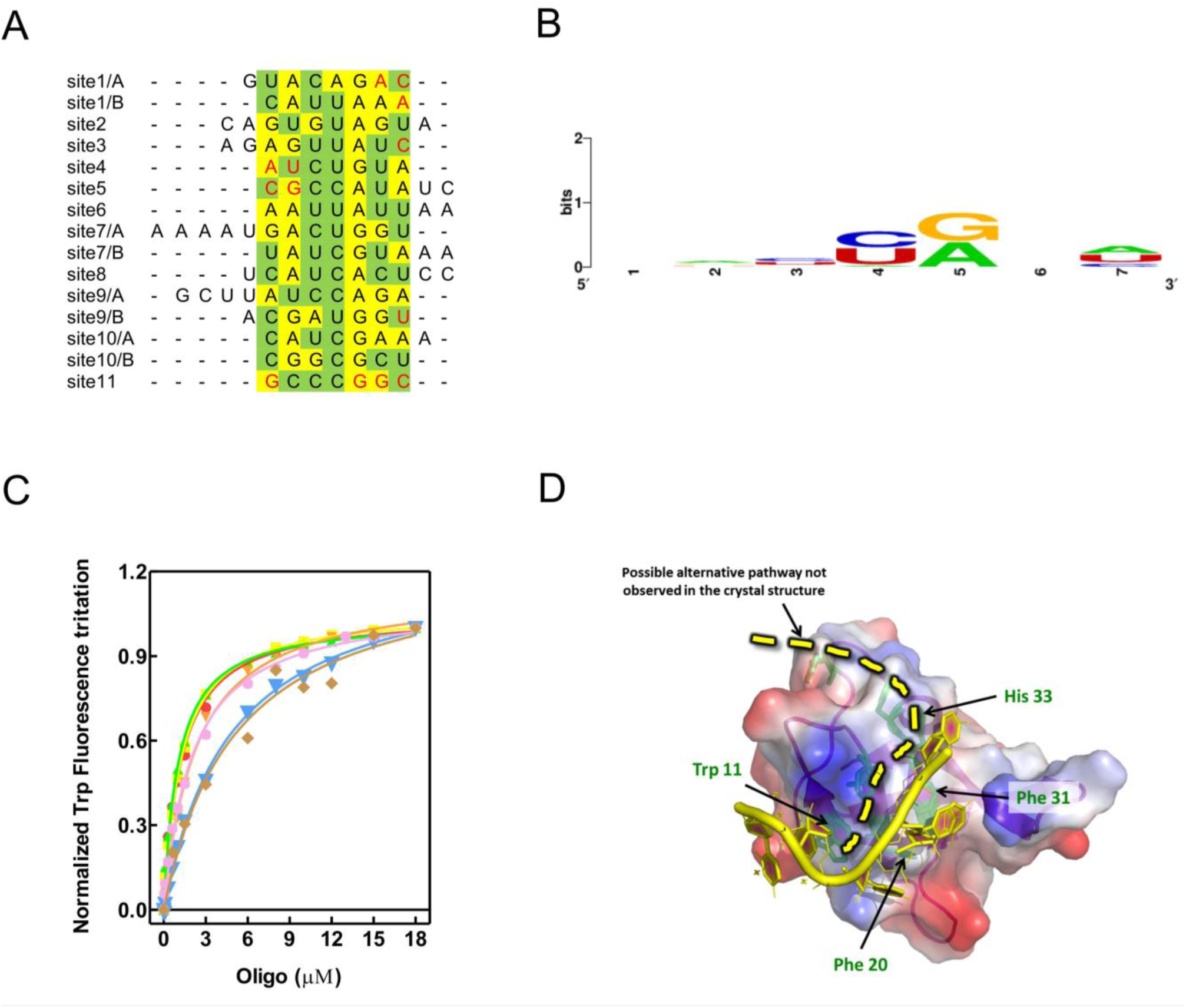
Characterization of CspA-RNA interaction. (A) Manual multiple alignment of the sites cross-linked/footprinted by CspA on *cspA* mRNA. Since CspB, the CspA-homologue of *Bacillus subtilis*, recognizes a sequence of 6-7 nts (Lopez and Makhatadze, 2000; Sachs et al., 2012), we hypothesized that the sites in which the cross-links/footprints were particularly extended (> 12 nts) could be the results of the binding of two adjacent CspA molecules. For this reason, we divided these extended sites (sites 1, 7 and 9) into two sub-sequences of comparable length, named A and B, which were used to build the alignment. The yellow and green highlighting indicates purines and pyrimidine, respectively. The bases in red are adjacent to those footprinted/cross-linked by CspA. (B) Logo representation of the CspA binding preference derived from the alignment shown in panel A and generated with WebLogo (Crooks et al., 2004). (C) Quenching of the CspA Trp fuorescence induced by the binding of oligo1 (5′-AACUGGUA-3′, red), which contains the most frequent bases found in the conserved positions of the Logo, or by the following RNA sequences: oligo2 (5′-AAAUGGUA-3′, orange); oligo3 (5′-AACAGGUA-3′, yellow); oligo4 (5′-AACUAGUA-3′, green); oligo5 (5′-AACUGAUA-3′, blue); oligo6 (5′-AACUGGAA-3′, pink) and oligo 7 (5′-AAAAAAAAA-3′, brown). Experiments were performed at 20°C in the presence of 1 µM of CspA and the indicated concentrations of oligos. Further details are given in Experimental procedures. (D) Model of CspA-oligoRNA interaction obtained using *B. subtilis* CspB-rC7 structure (pdb file 3pf4; Sachs et al., 2012).

**Table 1.** Equilibrium dissociation constants of the CspA:RNA oligonucleotide complexes determined by tryptophane fluorescence titration.

| Oligonucleotide | Sequence<br>5' to 3' | $K_D$, $\mu\text{M}$ |
| --- | --- | --- |
| Oligo 1 | AACUGGUA | $1.24 \pm 0.08$ |
| Oligo 2 | AAAUGGUA | $2.30 \pm 0.13$ |
| Oligo 3 | AACAGGUA | $1.24 \pm 0.05$ |
| Oligo 4 | AACUAGUA | $1.03 \pm 0.06$ |
| Oligo 5 | AACUGAUA | $4.86 \pm 0.24$ |
| Oligo 6 | AACUGGAA | $2.08 \pm 0.14$ |
| Oligo 7 | AAAAAAAAA | $5.13 \pm 1.15$ |

Because there is a high degree of sequence and structure similarity between *E. coli* CspA and its *B. subtilis* orthologue CspB, we produced a homology model of CspA in complex with Oligo 1 using the available 3D structure of a CspB-RNA complex (Sachs et al., 2012). This model (Fig. 9D) allowed us to verify that the YYR core motif, unlike other combinations of trinucleotides like RRR, fits very well in the binding pocket. From the model, the sidechains of His33, Phe31, Phe20, and Trp11, would stack with A2, C3, U4, G5, respectively, while G6 is stacked on G5. Hence, RNA binding is dominated by stacking interactions between the YYR motif and the aromatic protein sidechains of Phe31, Phe20 and Trp11. Furthermore, the purine downstream from the YYR motif can strengthen the stacking of the side chain of Trp11 with G5, while the nucleotide (A/U) upstream the core motif can stack with the sidechain of His31, further stabilizing the protein-RNA interaction.

Taken together, our data support that the YYR might be the preferred seed sequence to initiate binding. They are also in agreement with earlier works (Jiang et al., 1997; Lopez and Makhatadze, 2000) reporting the K_D_ for the CspA-RNA complex in the µM range.

### CspA promotes translation of numerous CS and non-CS mRNAs at low temperature

To gain additional insight into the CspA properties we tested the effect of CspA on the *in vitro* translation of mRNAs other than *cspA*. Two classes of mRNAs were selected to carry out this analysis: (i) the cold-shock transcripts *cspB* and *P1infA,* the former belonging to the *E. coli csp* gene family (Yamanaka et al., 1998) and the latter originating from the P1 promoter of *infA* and encoding Initiation Factor 1 (IF1) (Giangrossi et al., 2007); and (ii) the non-cold-shock transcripts *hupA* and *cspD*, encoding the α-subunit of the nucleoid associated protein HU (Giangrossi et al., 2002) and protein CspD (Yamanaka and Inouye, 1997), respectively. As shown in Fig. 10, the translation of *infA* and *cspD* mRNAs is strongly stimulated by CspA (3-4 fold), that of *cspB* mRNA is moderately enhanced, while *hupA* mRNA translation is insensitive to CspA addition. This result confirms that CspA is able to stimulate the translation at low temperature of various transcripts other than its own mRNA (Giuliodori et al., 2004), but also indicates that this activity cannot be generalized.

**FIGURE 10.**
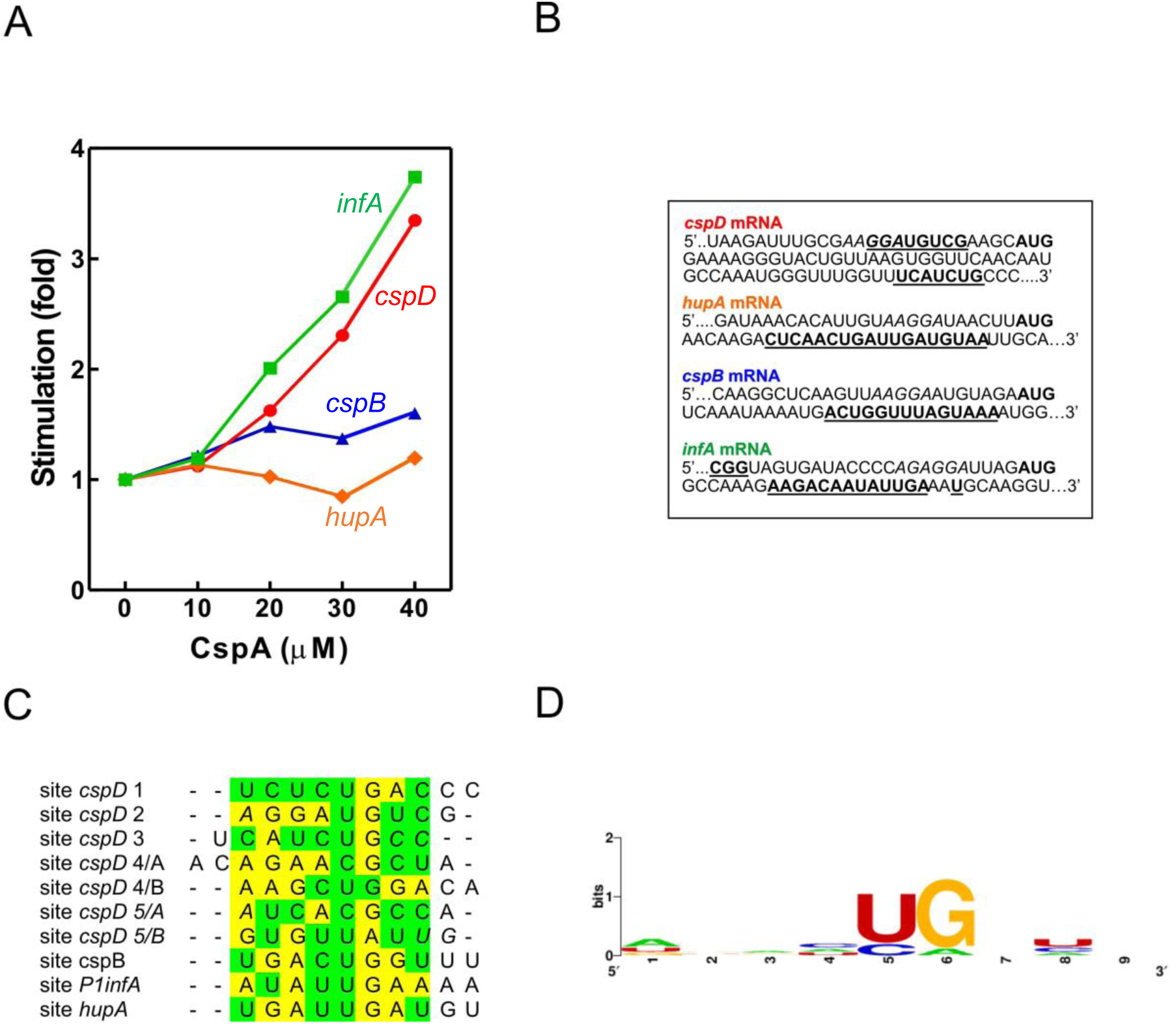
CspA binding to various mRNAs and functional effects. (A) *In vitro* translation at 15°C performed with control 70S and S100 in the presence of *P1infA* mRNA (green squares), *cspD* mRNA (red circles), *cspB* mRNA (blue triangles), *hupA* mRNA (orange diamonds) and the indicated amounts of purified CspA. (B) CspA binding sites (bold underlined) identified around the Translation Initiation Region (TIR) of the indicated mRNAs by crosslinking experiments performed at 15°C. SD sequence and start codon are indicated in italics and bold, respectively. (C) Multiple alignment of the sites crosslinked by CspA on the indicated mRNAs. As described in the legend of Fig. 9, we divided all sites made up of > 12 nts into two sub-sequences of comparable length, named A and B, which were used to build the alignment. The yellow and green highlighting indicates purines and pyrimidine, respectively. The bases in italics are adjacent to those crosslinked by CspA. (D) Logo representation of the CspA binding preference derived from the alignment shown in panel C and generated with WebLogo (Crooks et al., 2004).

The existence of the CspA-dependent translational stimulation of the tested mRNAs (*infA*, *cspD*, *cspB*) raises the question as to whether these transcripts were directly recognized by CspA. To address this issue, we mapped the possible interactions between CspA and our selected mRNAs by UV-induced crosslinking experiments at 15°C (Fig. 10B and Fig. 10-figure supplement 1). Notably, there appears to be a binding site common to all tested mRNAs – apart from *cspD* mRNA. The average length of the binding site is of 14 nts and is located between 9-12 nts downstream from the G of the translation initiation codon (Fig. 6, 7, 7 supplement 1, and 10 supplement 1). In the case of *cspD* mRNA, multiple cross-links were present along the entire mRNA (Fig. 10-figure supplement 1). In the region near the AUG codon, a cross-link of moderate intensity is observed in the SD region while a more intense one is located between the 17^th^ and the 19^th^ codons of the coding region. The YYR motif was found also in the regions cross-linked with CspA in these mRNAs. The multiple alignments built using the YYR motif as the reference sequence (Fig. 10C) produced a Logo similar to that generated using the binding sites on *cspA* mRNA (Fig. 10D).

## DISCUSSION

### The CspA paradox

Our data demonstrate that the binding of CspA to mRNA is not always accompanied with an effect on translation. This was particularly well illustrated with *cspA* mRNA: despite the extensive binding of CspA to the cold-form of *cspA* mRNA, translation of this structure is not stimulated, whereas the translation of the 37°C-form is enhanced by CspA although CspA binding is less efficient. How can this apparent paradox be explained?

Immediately after cold-shock CspA becomes a very abundant protein (Brandi et al.,1999), and it is estimated to be bound in several copies to cellular mRNAs (Ermolenko and Makhatadze, 2002). CspA was shown to bind its own mRNA (Jiang et a., 1997) and to act as an RNA chaperone (Jiang et a., 1997; Rennella, 2017; Zhang et al., 2018). In this work, we demonstrated that CspA is able to recognize in its mRNA short and degenerated sequences mostly located in single stranded regions, including internal and apical loops. Furthermore, we showed that CspA stimulates the translation at low temperature from its 37°C-form mRNA, which adopts a large and irregular hairpin structure sequestrating the SD sequence (Fig. 7). This CspA-dependent translational stimulation is observed with other mRNAs. In all tested mRNAs, with the exception of *cspD*, we identified a cross-link positioned between 9-12 nts from the initiation triplet. We propose that this region of mRNAs could be a preferential CspA binding site since it is usually poorly structured (Del Campo, 2015). In spite of this interaction, our results demonstrate that CspA enhances the translation of only some of the mRNAs that it is able to bind. Our functional experiments performed with *cspA* mRNA demonstrate that the translational stimulation affects elongation rather than initiation. Particularly, the RelE-walking experiment proves that this activity consists in facilitating ribosome progression on the mRNA at low temperature.

During translation the ribosome is able to melt secondary structures of the mRNA thanks to the helicase activity of S3, S4 and S5 proteins (Qu et al., 2011). It is very likely that this activity could be partly impaired by the low temperature, which is known to stabilize base pairing interactions, making it harder for the ribosome to melt the secondary structures (Liu et al., 2014). The presence of CspA on the mRNA could be useful to facilitate ribosome progression either by destabilizing some positions and/or by preventing the re-formation of the secondary structures after the first elongating ribosome has unwound them.

Based on our data, two mechanistic models, not mutually exclusive, can co-exist. The first model builds on the CspA RNA chaperone activity (Rennella et al., 2017). This model outlines how CspA stimulates annealing of two complementary RNA hairpins by weakening the RNA base pairing interactions which prevent the RNA to reach its final and less energetic state, to form a thermodynamically more stable hetero-duplex. Therefore, in the presence of translating ribosomes, the destabilization of mRNA structures will increase the proportion of single-stranded regions thus facilitating ribosome progression. In the second model, the amount of the mRNA-bound CspA increases as the translating ribosomes open up the mRNA structures. In this case, stimulation during translation would not depend on the amount of CspA pre-bound to the mRNA but rather on the capacity of CspA to rapidly bind (or re-bind) the regions melted by the passage of the first ribosome and keep them single-stranded. This RNA chaperone activity has been dubbed “overcrowding” (Cristofari and Darlix, 2002).

Both models would predict an easy displacement of CspA molecules as the ribosomes transit on the mRNA interaction sites and can explain the “CspA paradox” (Fig. 11). In fact, ribosome progression would be stimulated by CspA only with structured mRNAs whose conformational state is stabilized by the low temperature, while the effect will not be seen with mRNAs carrying a more open conformation, intrinsically suitable for translation at low temperature. It can thus be predicted that translation of mRNAs that are too structured and that contain too few sites for CspA binding would be little stimulated by CspA.

**FIGURE 11.**
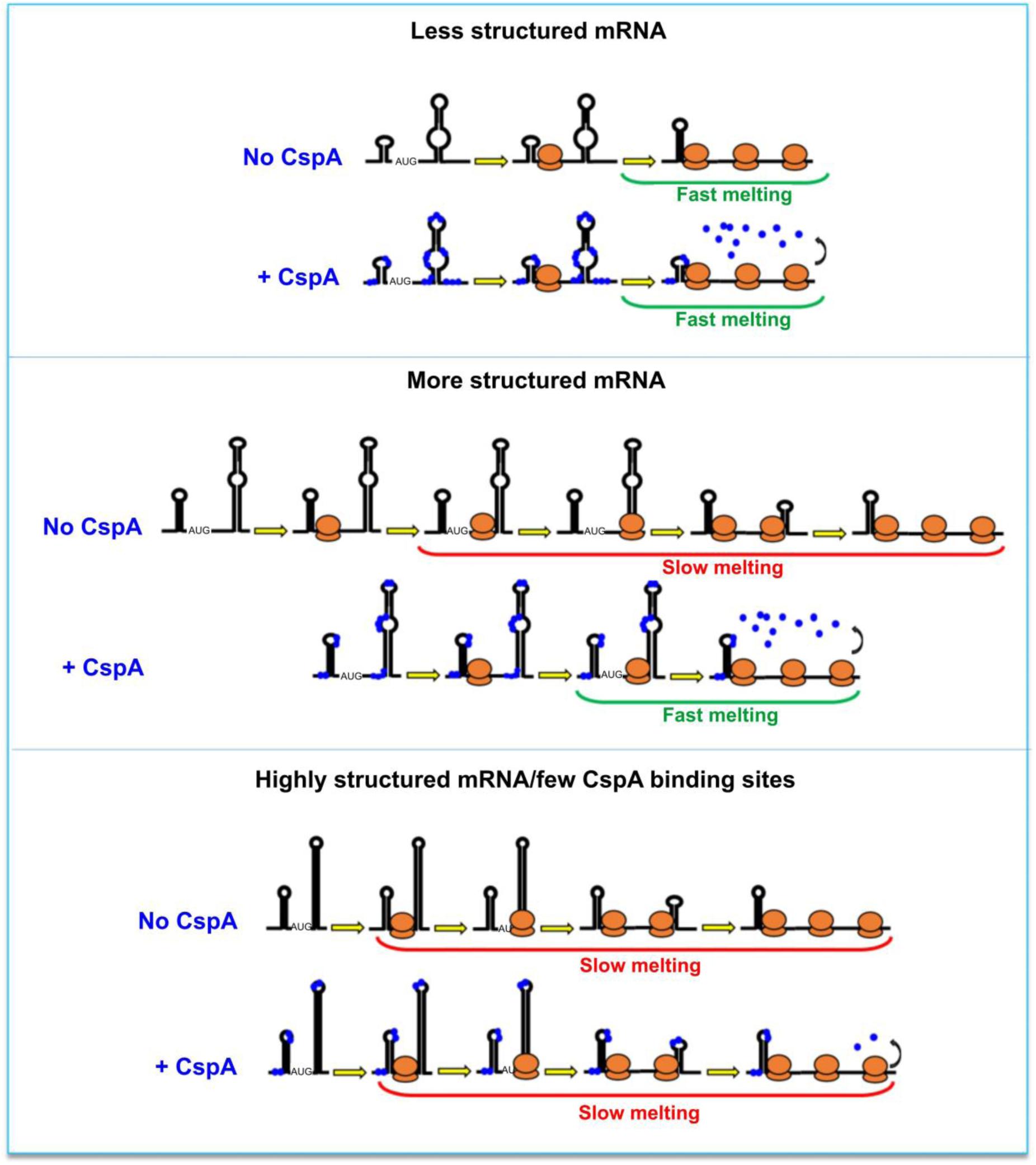
The CspA paradox. CspA would ensure ribosome progression by maintaining the bound regions unstructured and/or destabilizing the helices when ribosomes translate along the mRNA. This effect will not be seen with mRNAs attaining an open conformation compatible with translation at low temperature, or in the case of mRNAs that are either structured or containing few sites suitable for CspA binding.

### Stimulation of cspA mRNA translation is one of the first tasks of CspA during cold acclimation

At the beginning of the acclimation phase, both CspA and its mRNA are already present and the translation activation could be rapidly achieved through their interaction. Indeed, immediately after the cold stress the *cspA* mRNA transcribed at 37°C is stabilized 100 times, with its half-life increasing from 12 seconds to 20 minutes (Fang et al., 1997; Goldenberg et al., 1996). CspA protein is also known to be abundant at 37°C during early exponential growth, when its concentration reaches up to 50 µM (Brandi et al., 1999). Under these conditions the cold-shock induction of *cspA* is rather low (only about 3-fold) and at the onset of the cold stress the cells use predominantly *cspA* mRNA and CspA protein already synthetized at 37°C. Therefore, the capacity of CspA to favor the translation of its mRNA folded in the 37°C-conformation in the cold can speed up the accumulation of more *c*spA product, in a positive auto-regulatory loop. Our data suggest that the translational stimulation by CspA should take place at concentrations ≥ 25 µM. In addition, probing/footprinting experiments suggested that CspA does not produce important conformational changes on the full-length pre-folded *cspA* mRNA, even at concentrations > 100 µM. However, CspA seems to affect the conformation of a *cspA* mRNA fragment corresponding to the first 87 nts at concentrations > 50 µM. Therefore, CspA could play different roles in the cell depending on its expression level. When present at concentrations < 50 µM, its main effect would be to favor the progression of the ribosome on its and other mRNAs, whereas at higher concentration, CspA could have an impact on the co-transcriptional folding process of its targets, *cspA* mRNA *in primis*. This hypothesis is supported by the recent work of Zhang and coworkers (2018), which have demonstrated that at a concentration of 100 µM, CspA can modulate the structure of its mRNA, as well as that of *cspB*, thereby making it more susceptible to degradation at the end of the acclimation phase. Zhang *et al*. have also shown that CspA contributes to support translation recovery of the other genes during cold-shock. This result is in agreement with our data, which show that CspA favors the translation of other mRNAs. Therefore, we have re-analyzed ribosome profiling data from Zhang *et al*. to understand if we could observe *in vivo* the same mechanism of stimulation of translation progression from initiation to elongation operated by CspA. Analogously to our *in vitro* RelE walking assay, ribosome profiling experiments allow to map ribosome pausing or stalling sites, which are evinced by peaks of ribosome protected fragments. Interestingly, in the cell subjected to cold shock the peaks at the initiation codons of the mRNAs tested in this *as well as in a previous work* (Giuliodori et al., 2004) progressively decrease with the accumulation of CspA during the acclimation phase (Fig. 12). These data confirm the possibility that our proposed model for CspA stimulation of translation could take place *in vivo* on different mRNAs.

**FIGURE 12.**
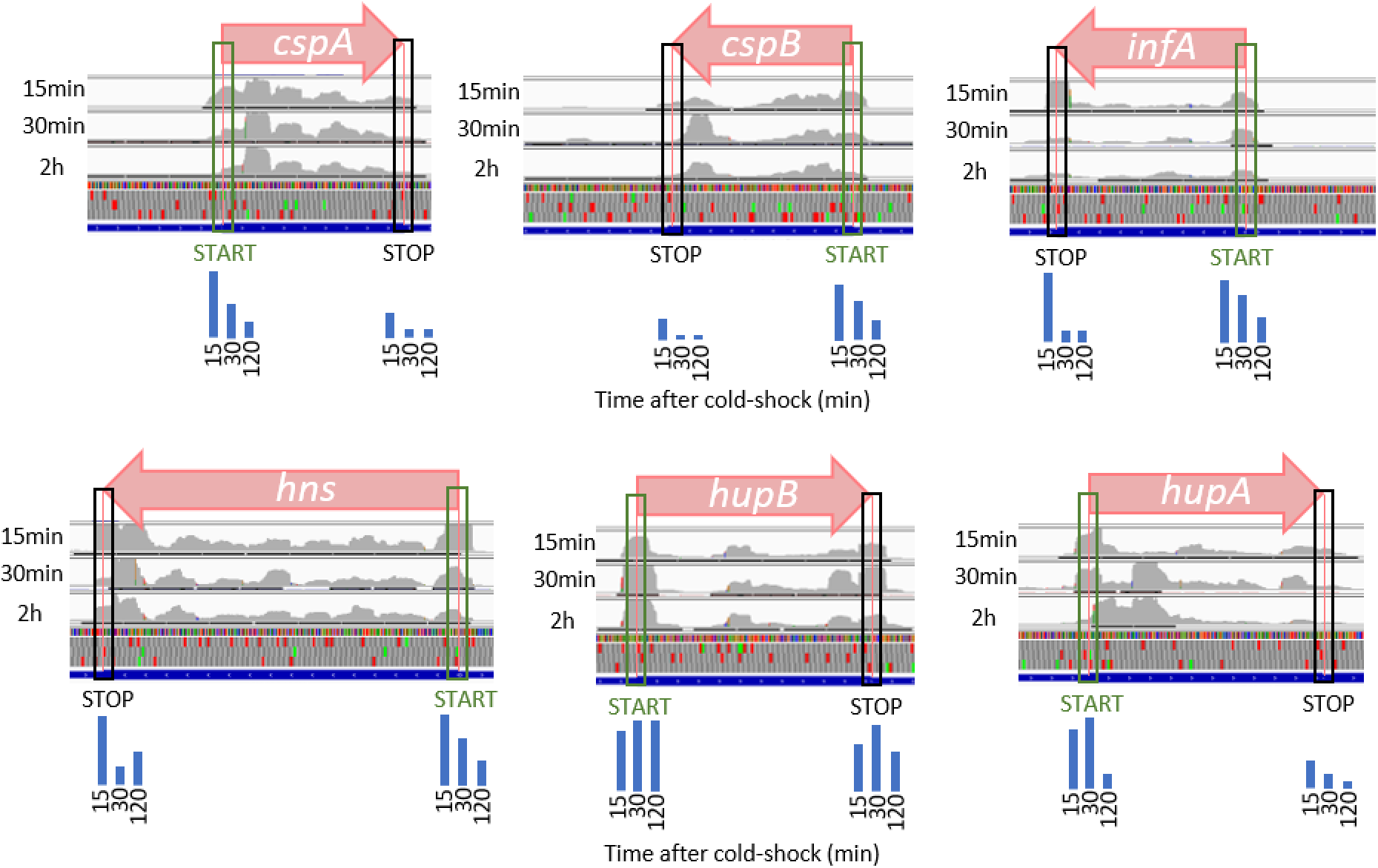
*In vivo* analysis of translation progression during cold acclimation. Ribosome profiling data obtained at 15 minutes, 30 minutes and 2 hours after temperature down-shift (10 °C), have been obtained from GEO series GSE103421 (Zhang et al., 2018). Row data have been trimmed of adapter sequences and bad quality reads, before being aligned on *E. coli* genome (NC_000913.3). Coverage tracks of aligned reads from ribosome protected fragments indicate the depth of the reads displayed at each locus. To compare different positions of the same region (coding sequence), each track has been normalized on the average of the density peaks of the region, excluding the peaks on the start and stop codons. The blue bars represent the quantization of peak densities at the start and stop codons at the three time points. From left to right, 15 minutes, 30 minutes and 2 hours, respectively. Normalized coverage tracks for *cspA, cspB, infA* and *hns* mRNAs show peaks on start codons which progressively decrease during cold acclimation. Normalized coverage tracks for *hupA* and *hupB* mRNAs do not show the same trend and their initiation peaks remain quite high even after several minutes of cold acclimation. *CspD* mRNA is not expressed under these conditions.

In addition to CspA orthologues present in all bacterial taxa, multiple paralogues have wide evolutionary distribution (Graumann and Marahiel, 1996). Some of these paralogues carry out overlapping functions, as demonstrated by the fact that in *E. coli* four out of nine *csp* genes must be deleted to obtain a cold-sensitive phenotype and that the overexpression of any member of the *csp* family (except for *cspD*) suppresses the phenotype (Xia et al., 2001). It is also known that these small proteins act as anti-termination factors during transcription and can bind both ssDNA and RNA with different specificity. CspB, CspC and CspE display specificity for 5′-UUUUU-3′, 5′-AGGGAGGGA-3′ and 5′-AAAUUU-3′ sequences, respectively, with K_D_ values in the range of 1-10 µM, similar to that calculated for CspA. Thus, it is conceivable that during cold-shock various Csp could be bound to the mRNAs to regulate their transcription or modulate their structure, stability and translation, possibly in a concentration-dependent manner as in the case of CspA.

## MATERIALS AND METHODS

### General preparations and buffers

*Escherichia coli* MRE600 70S ribosomes, S100 post-ribosomal supernatant, 30S ribosomal subunits and purified initiation factors IF1, IF2, and IF3 were prepared as described previously (Giuliodori et al., 2004; Giuliodori et al., 2007).

The following buffers were used:

*Buffer A*: 25 mM tris-HCl, pH 8.5, 5% glycerol, 100 mM NaCl, 0.025% Nonidet P40; *Buffer B*: 25 mM tris-HCl, pH 8, 1.3 M NaCl, 5% glycerol, 6 mM β-mercaptoethanol, 0.1 mM PMSF, 0.1 mM benzamidine; *Buffer C*: 25 mM tris-HCl, pH 8.0, 700 mM NaCl, 5% glycerol, 6 mM β-mercapto-ethanol, 0.1 mM PMSF, 0.1 mM benzamidine; *Buffer D*: 25 mM tris-HCl, pH 8.0, 300 mM of NaCl, 5% glycerol, 20 mM Imidazole, 6 mM β-mercaptoethanol, 0.1 mM PMSF, 0.1 mM benzamidine; *Buffer E*: 25 mM tris-HCl, pH 8.0, 300 mM of NaCl, 5% glycerol, 300 mM Imidazole, 6 mM β-mercaptoethanol, 0.1 mM PMSF, 0.1 mM benzamidine; *Buffer F*: 25 mM tris-HCl, pH 8.0, 100 mM NaCl, 5%glycerol, 6 mM β- mercaptoethanol, 0.1 mM PMSF, 0.1 mM benzamidine; *Buffer G*: 25 mM tris-HCl, pH 8.0, 300 mM NaCl, 5% glycerol, 6 mM β-mercaptoethanol, 0.1 mM PMSF, 0.1 mM benzamidine; *Buffer H*: 20 mM tris-HCl, pH 7.1, 10 mM NH_4_Cl, 1 mM MgCl_2_, 10% glycerol, 0.1 mM EDTA, 6 mM β-mercaptoethanol; *Buffer I*: 20 mM Hepes-KOH, pH 7.5, 10 mM MgCl_2_, 50 mM KCl; *Buffer L*: 10 mM Tris-HCl pH 7.5, 60 mM NH_4_Cl, 1 mM DTT, 7 mM MgCl_2_; *Buffer M*: 20 mM Tris-HCl pH 7.5, 60 mM KCl, 1 mM DTT, 10 mM MgCl_2_; *Buffer N*: 20 mM Na-cacodylate, pH 7.2, 10 mM MgCl_2_, 50 mM KCl; *Buffer O*: 20 mM Tris-HCl pH 7.5, 60 mM KCl, 40 mM NH_4_Cl, 3 mM DTT, 10 mM MgCl_2_, 0.002mg/ml BSA; ITC buffer: 20 mM Tris-HCl pH 7.1, 10 mM NH_4_Cl, 7 mM MgCl_2_, 10% glycerol, 0.1 mM EDTA, 6 mM β-mercaptoethanol.

### Molecular cloning, expression and purification of CspA

The coding region of *E. coli cspA* gene was amplified by PCR from the pUT7cspA construct (Giuliodori et al., 2010) using the forward primer G655 5′-CATGCCATGGCCGGTAAAATGACTGGTATCG-3′ and the reverse primer G656 5′-CGGGATCCTTACAGGCTGGTTACGTTAC-3′ and cloned into the pETM11 vector (Dümmler et al. 2005) using NcoI and BamHI restriction sites. Because the introduction of the NcoI restriction site had changed the second amino acid of the CspA sequence, after the molecular cloning the wt sequence was restored by mutagenesis using the QuikChange Site-Directed Mutagenesis Kit (Agilent Technologies, Inc., Santa Clara, CA), the pETM11-CspA plasmid as DNA template and the mutagenic primers G670 5′-TTTCAGGGCGCCATGTCCGGTAAAATGACTG-3′ and G671 5′-CAGTCATTTTACCGGACATGGCGCCCTGAAA-3′.

Overproduction of protein CspA was induced into a culture of *E. coli* BL21 (DE3)/pLysS cells grown in LB medium at 37°C till OD_600_ = 0.4 by the addition of 1 mM of isopropylbeta-D-1-thiogalactopyranoside (IPTG). After transferring the culture to 20°C for 12 h, cells were harvested by centrifugation and the pellet resuspended in Buffer A and stored at - 80°C. After thawing, cells were diluted in an equal volume of Buffer B and lysed by sonication. The resulting cell extract, cleared by centrifugation, was loaded onto a nickel-nitrilotriacetic acid (Ni-NTA) chromatographic column equilibrated in Buffer C. After washing in Buffer D, protein CspA was eluted using Buffer E, pooled and dialyzed against Buffer F. To remove the His-Tag sequence, 15 mg of CspA were incubated for 4 h at 20°C with the His-Tag TEV protease (Kapust et al. 2002). At the end of the incubation, the concentration of NaCl was increased to 300 mM and the cleaved CspA was loaded onto a Ni-NTA column equilibrated in Buffer G. The flow-through, containing CspA with no His-Tag, was dialysed overnight at 4°C against Buffer H. Then, CspA was concentrated by centrifugation in Microcon tubes (Amicon-Millipore) with 3 KDa cut-off at 13.8 krcf, 4°C, until the concentration was ≥ 400 µM and stored at -80°C in small aliquots. The purity of CspA protein was checked by 18% SDS-PAGE.

### mRNA preparation

The DNA templates used *for in vitro* transcription of the various mRNAs were constructed as specified in Giuliodori et al., 2010 and Di Pietro et al., 2013. All mRNAs obtained by *in vitro* transcription with T7 RNA polymerase were purified and labelled as described in Giuliodori et al., 2010.

### Translation assays

Before use, the mRNAs were denatured at 90°C for 1 min in RNase free H_2_O and renatured for 15 min at 15°C or 37°C in Buffer I. When required, CspA was added after renaturation at the concentrations indicated in the figures.

*In vitro* translation reactions were carried out in 30 µL containing 20 mM Tris-HCl, pH 7.7, 12 mM Mg acetate, 80 mM NH_4_Cl, 2 mM DTT, 2 mM ATP, 0.4 mM GTP, 10 mM phosphoenolpyruvate, 0.025 µg of pyruvate kinase/µL reaction, 200 µM of each amino acid (minus Alanine), 5 µM [^3^H] Alanine (309 mCi/mmol), 50 mM cold Alanine, 0.12 mM citrovorum (Serva) and 0.4 U/µL of RNasin (Promega). The reaction mixture also contained 30 pmoles of *in vitro* transcribed mRNAs and either the amount of S30 crude extracts corresponding to 20 pmoles of 70S ribosomes or 30 pmoles of purified 70S ribosomes, 15 pmoles of purified Initiation Factors IF1, IF2 and IF3, and 2 µL of S100 post-ribosomal supernatant. After incubation for the indicated times and temperatures, samples (15 µL) were withdrawn from each reaction mixture and the incorporated radioactivity determined by hot-trichloroacetic acid (TCA) method.

Initiation complex (IC) formation assays (filter binding) were carried out in 30 µL of Buffer L using 0.5 µM 30S ribosomal subunits either alone (for the 30S IC) or in the presence of 1 µM of 50S subunits (for the 70S IC), 0.5 µM ^35^S-fMet-tRNA, 0.5 mM GTP, 0.5 µM IF1, 0.5 µM IF2, 0.5 µM IF3, 1 µM *cspA* and 0.4 U/ µL of RNasin (Promega). Binding of ^32^P-labelled *cspA* mRNA to 30S subunits was performed in 40 µl of Buffer L containing 20 pmoles of *cspA* mRNA and 9000 cpm of [^32^P] *cspA* mRNA, 0.4 U/µL of RNasin (Promega), 20 pmoles of 30S subunits and either 30 pmoles of tRNA_i_ or 30 pmoles of fMet-tRNA_i_ and 20 pmoles of IF1, IF2 and IF3. After 30 min incubation at 15°C, the amount of initiation complex formed was determined either by filtration through 96-multiscreen-HTS-HA Millipore plates (mRNA binding) or by nitrocellulose filtration (30S and 70S IC), followed by liquid scintillation counting.

The toeprinting assay was performed essentially as described (Fechter, et al, 2009). The reaction was carried out in 10 μl of Buffer M containing 0.4 U/µL of RNasin (Promega) in the presence of 0.02 μM *cspA* mRNA, 4 μM tRNA_i_, 50 μM each of dNTPs, P-labeled oligo csp2 (5′-CGAACACATCTTTAGAGCCAT-3′), and 0.2 μM of *E. coli* 30S subunits. The reaction mixtures were incubated for 30 min at 15°C. Primer extension was conducted with 4 units of Avian Myeloblastosis Virus (AMV) reverse transcriptase (Sigma) for 1 hour at 15°C. The reaction products were analysed on 8% PAGE-urea gel.

### RNA footprinting assays

Before use, the RNAs were denatured at 90°C for 1 min in RNase-free H_2_O and renatured for 15 min at 15°C or 37°C in the buffers used for enzymatic probing or hydroxyl radical cleavage experiments.

Enzymatic probing was carried out on ^32^P-end-labeled transcripts (50,000 cpm) essentially as described earlier (Giuliodori et al., 2010) after incubating the renatured mRNA with the amounts of CspA indicated in the figure legends.

Probing by hydroxyl radical cleavage was performed essentially as described (Fabbretti et al., 2007). CspA was allowed to bind *cspA* mRNA in 40 µL of Buffer N by incubating 10 pmoles of renatured mRNA with the indicated amounts of protein CspA in the presence of 0.4 U/µL of RNasin (Promega). After 15 min at 15°C, H_2_O_2_ was added (0.15% final concentration) and the cleavage started by adding Fe(II)-EDTA (3 mM final concentration). Cleavage was allowed to proceed for 15 sec at 15°C before addition of 260 µl quenching solution containing 0.3 M Na acetate (pH 5.2) in absolute ethanol. The precipitated samples were resuspended in H_2_O, extracted with phenol-chloroform and re-precipitated with cold 0.3 M Na acetate (pH 5.2) in absolute ethanol. These reaction products, resuspended in 3 µL of sterile H_2_O, were then subjected to primer extension analysis as described earlier (Fabbretti et al. 2016) using cspA1, csp2 and cspA3 primers (Giuliodori et al., 2010).

### CspA-RNA cross-linking

For the CspA-RNA cross-linking experiments, 0.02 µM of the ^32^P-labeled primers indicated in the figure legends were mixed with 0.35 µM of the corresponding mRNAs. After a denaturation step at 90°C for 1 min, the samples were incubated at either 15°C or 37°C for 10 min in Buffer I containing 0.4 U/µL of RNasin (Promega). Following renaturation, the reaction mixtures were dispensed in tubes containing increasing amounts of purified CspA (reaction volumes: 10 µL) and the protein was allowed to bind for 10 min at the indicated temperatures. Subsequently, the samples were transferred to an ice-cold plate and U.V. irradiated for 2 min using the GS Gene-linker BioRad (180 mJ, 254 nm bulbs at 12 cm from the U.V. source). The cross-linked RNA was primer-extended using AMV Reverse Transcriptase as previously described (Giuliodori et al., 2010).

### Isothermal titration calorimetry (ITC)

All samples were dialyzed against ITC buffer using centrifugal filter units (Centricon, Merck Millipore), 3 K for CspA and 100 K for *E. coli* ribosome. ITC experiments were done on the microcalorimeter MicroCal PEAQ-ITC (Microcal-Malvern Panalytical, Malvern, UK). For CspA/ribosome binding studies, experiments were done by successive injections of CspA in 30S, 50S or 70S solution at three different temperatures (15, 25 and 35°C). Data were analyzed with MicroCal PEAQ-ITC Analysis Software.

### RelE walking assay

RelE toxin was expressed and purified as described earlier (Andreev et al. RNA 2008). Ribosome progression on *cspA* mRNA was monitored by analysing the amount of RelE cleavages obtained by ribosomes paused at different sites on the mRNA. *In vitro* translation was carried out in 10 µL using the PURExpress kit (NEB) according to the commercial protocol in the presence of a mix of ^32^P-radiolabeled *cspA* mRNA (200000 cpm/µL) and cold *cspA* mRNA (0.8 µM), previously folded at 15°C or 37°C. When present, CspA was added at concentration of 30 µM. The reaction was incubated 2h at 15°C, and blocked by addition of chloramphenicol (1 mM) and different concentration of RelE as in figure 2, for 15 min at 15°C. The RNA fragments were then phenol extracted, subjected to 8% PAGE-urea and revealed by autoradiography. Quantization of each band was done using ImageQuant TL (GE Healthcare) and signal normalization was done using the sum of the quantization of all the bands present in each lane.

### RNA Electrophoretic mobility shift assay

Radiolabelled purified *187cspA* RNA (Giuliodori et al., 2010), 50000 cps/sample, at concentration < 1 pM, was denatured and renaturated at 15°C or 37°C, as described above. For each experiment, increasing concentrations of purified CspA (30-211 µM) were added to the 5′ end labelled *187cspA* RNA in a total volume of 10 µL in Buffer O containing 0.4 U/µL of RNasin (Promega). Complex formation was performed at 15°C or 37°C for 15 min. After incubation, 10 µL of glycerol blue was added and the samples were loaded on a 10% PAGE under non-denaturing conditions (1h, 300 V, 4°C).

### Steady-state fluorescence spectroscopy

To measure binding affinity between CspA and different RNA oligonucleotides, intrinsic tryptophan fluorescence quenching experiments with 1 μM of CspA and increasing amount of RNA oligonucleotides was performed in Buffer H. Fluorescence measurements were performed in quartz cells at 20 ± 0.5°C on a Fluoromax-4 fluorimeter (HORIBA Jobin-Yvon Inc., NJ., USA). The excitation wavelength was set at 295 nm for selective excitation of tryptophan residues and the emission wavelength was scanned from 305 to 450 nm. The peak of emission at 351 nm was used to measure the quenching effect. Increasing amounts of RNA (from 0.05 μM to 18 μM as described in figure 6) were added and the quartz cell was rapidly homogenized before fluorescence emission measurements. Fluorescence intensities were corrected for buffer fluorescence and dilution effects. Binding parameters were calculated as described in Dubois et al., 2018.

### 3D model of CspA-target RNA interaction

CspA crystal structure (pdb file 1MJC (Schindelin et al., 1994)) was superposed to the crystal structure of *B. subtilis* CspB in complex with an eptanucleotide RNA oligo (GUCUUUA) (pdb file 3PF4 Sachs et al., 2012). Oligo 1 (AACUGGUA) sequence was then modelled on the RNA structure by Assemble2 software (Jossinet et al., 2010).

## ACKNOWLEDGEMENTS

We thank Claudio Gualerzi, Pascale Romby and Serena Bernacchi for critical reading of the manuscript and helpful discussions. We thank Benoît Meyer for the help in the ITC experiments.

## Author contributions

Conceptualization, A.M.G. and S.M.; Methodology, A.M.G and S.M; Investigation, A.M.G., M. D., R. B., R. G., E. S. and E. E.; Writing – Original Draft, A.M.G. and S.M.; Writing – Review & Editing, A.M.G., M. D., R. B., R. G., E. S., V. H., E.E. and S.M; Visualization, A.M.G.; Supervision, A.M.G., E.E. and S.M.; Resources, A.M.G., E.E. and S.M.

patent US7118883b2), registrato 23 ottobre 2001, Inoue, Ueda.

---

**FIGURE 4-supplement 1.**
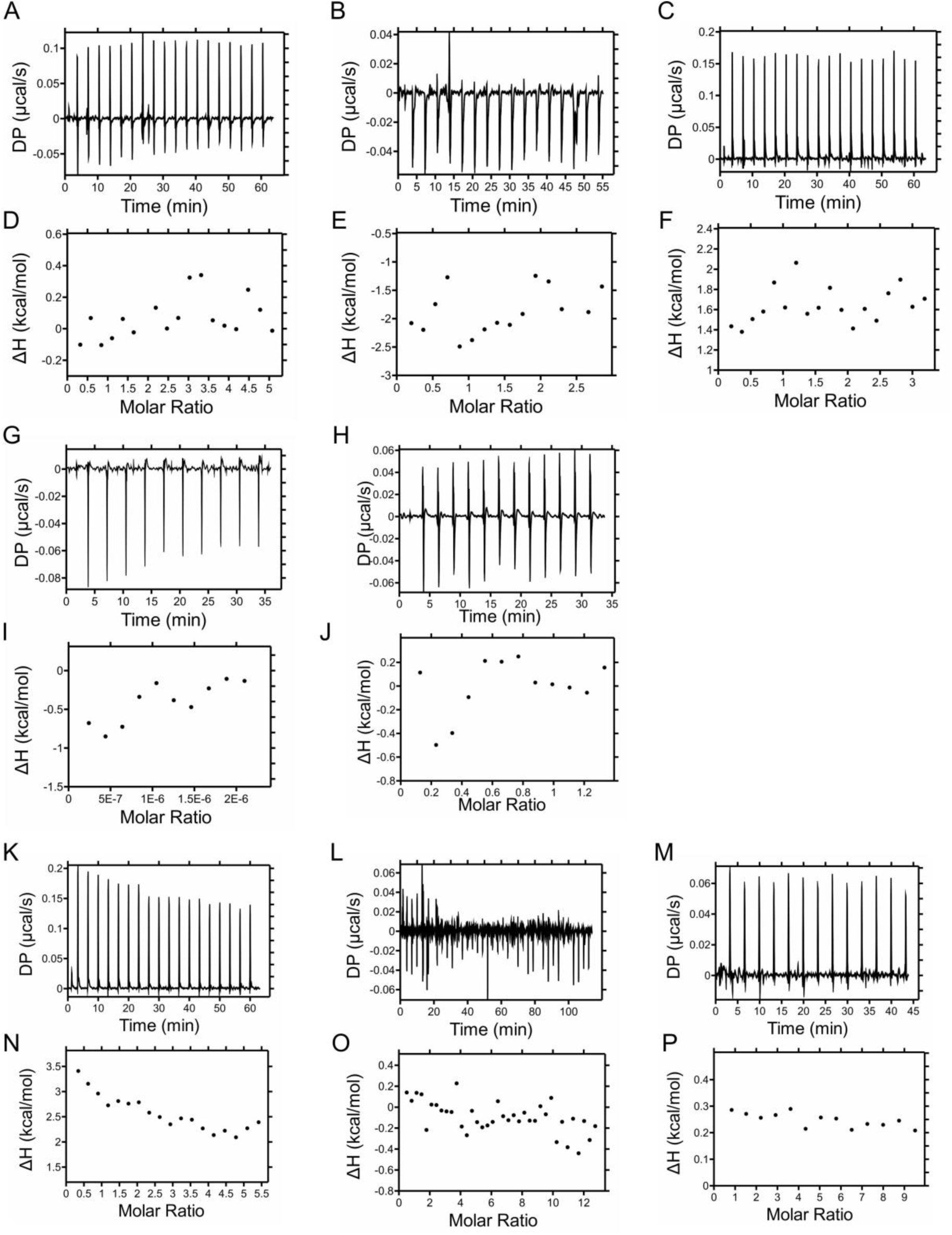
Binding of CspA to 70S ribosomes or ribosomal subunits. Titration of *E. coli* ribosomal particles with CspA was studied at 25°C (A-E) and 35 °C (F-H). Titration was carried out by consecutive 2 µL injections of CspA in a cell containing 200 µL of either 70S ribosomes (A, and F), 30S subunits (B and G) or 50S subunits (C and H). D and E are examples of signals of CspA and *E. coli* 50S subunits obtained upon dilution in ITC buffer at 25 °C: in (D) 2 µL of CspA were repeatedly injected in the sample cell filled with ITC buffer, while in (E) 2 µL of ITC buffer were repetitively injected in the sample cell filled with 50S subunits.

**FIGURE 7-supplement 1.**
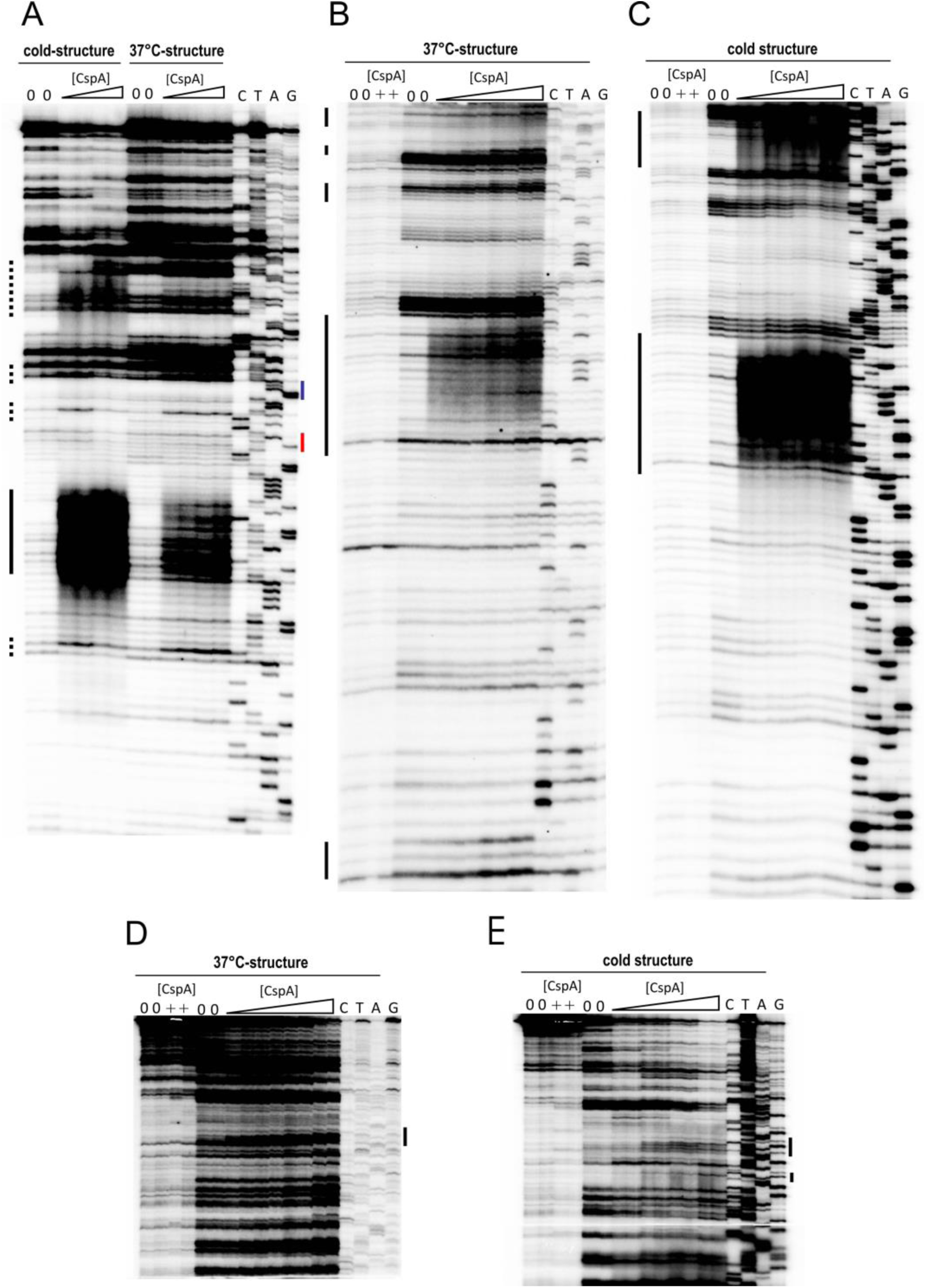
Localization of CspA binding sites on *cspA* mRNA by UV-induced cross-linking at 15°C. The experiments were performed in the absence (lanes 0) or in the presence of 28 and 57 µM of CspA (panel A) or 28, 57, 115, 170 µM of CspA (panels B, C, D, and E) with *cspA* mRNA folded in the conformations indicated at the top of the gels. Lanes C: mRNA alone, no cross-links; lanes +: mRNA+170 µM of CspA, no cross-links. Increasing concentrations are indicated with a triangle. All conditions are in duplicate. Lanes C, T, A, and G correspond to sequencing reactions. Primer extension analysis was performed using primer csp2 (panel A), which annealed to the coding region and primer csp3 (panels B, C), which annealed to the 3′ UTR. Panels D and E show the upper parts of the gels displayed in panels B and C. Bases whose accessibility to the cleavage is affected by CspA are highlighted by black bars on the left side of the gels. The blue and red bars on the right side of panel A indicate the Shine-Dalgarno sequence and the AUG initiation triplet, respectively.

**FIGURE7-supplement 2.**
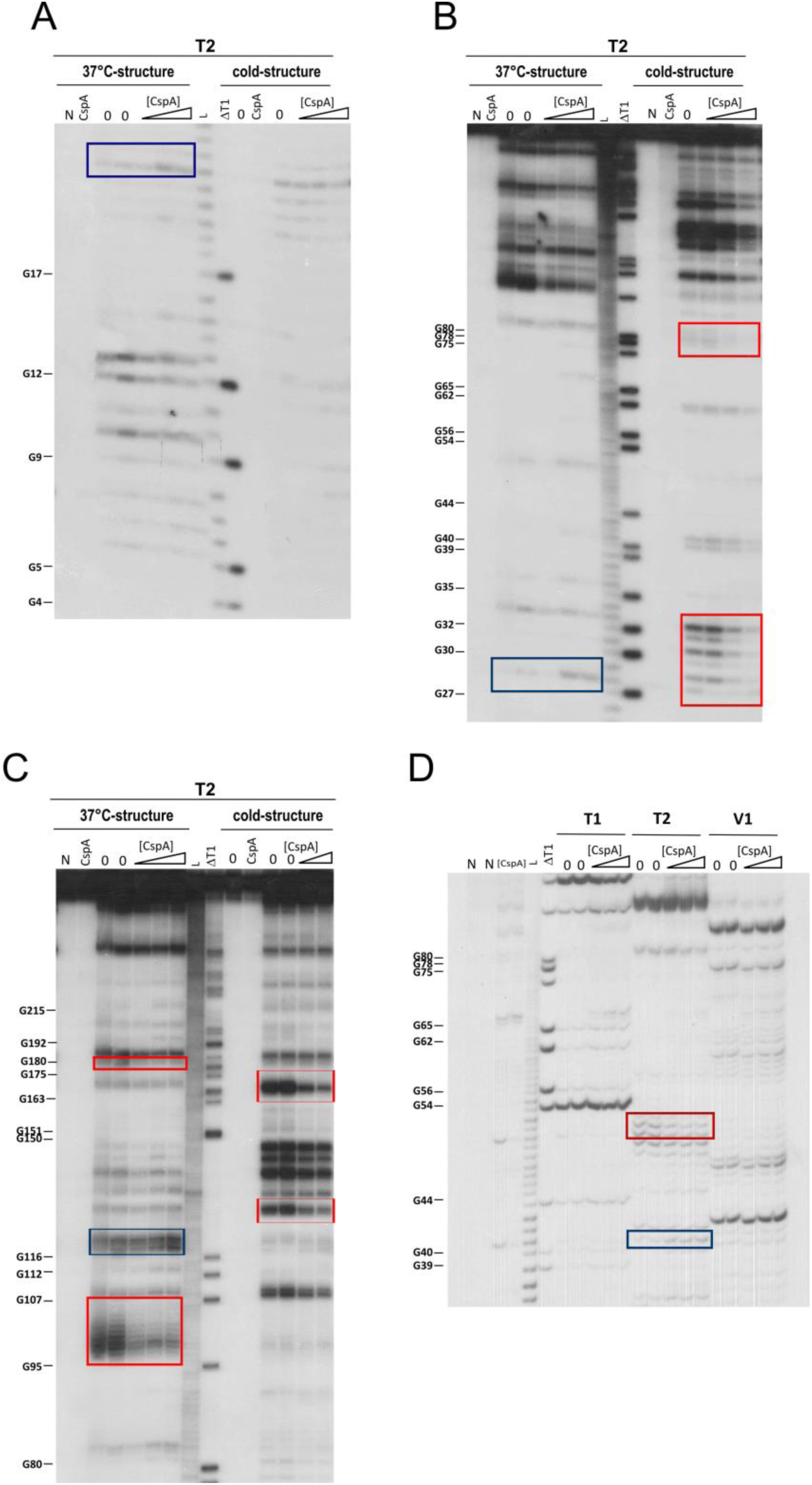
Footprinting experiments using RNases. Short (A and B) and long (C) electrophoretic migration of the fragments generated by RNase T2 (T2) digestion of 5′-end [^32^P]-labelled *cspA* mRNA folded in the conformations indicated at the top of the autoradiographies; (A) bottom and (B) top of the gel. (D) Short electrophoretic migration of the fragments generated by RNases T1, T2, and V1 digestion of 5′-end [^32^P]-labelled *cspA* mRNA folded in the 37°C- conformation. The experiment was carried out at 15°C in the absence (lanes 0) or in the presence of 80, 102, 160 µM of CspA (increasing concentrations are indicated with a triangle). Lanes N: controls with neither T2 nor CspA; lanes +: controls without T2, with CspA; lanes T: RNase T1 cleavages under denaturing conditions; lanes L: alkaline ladder. The red and blue boxes indicate the positions protected or exposed by CspA, respectively.

**FIGURE 7-supplement 3.**
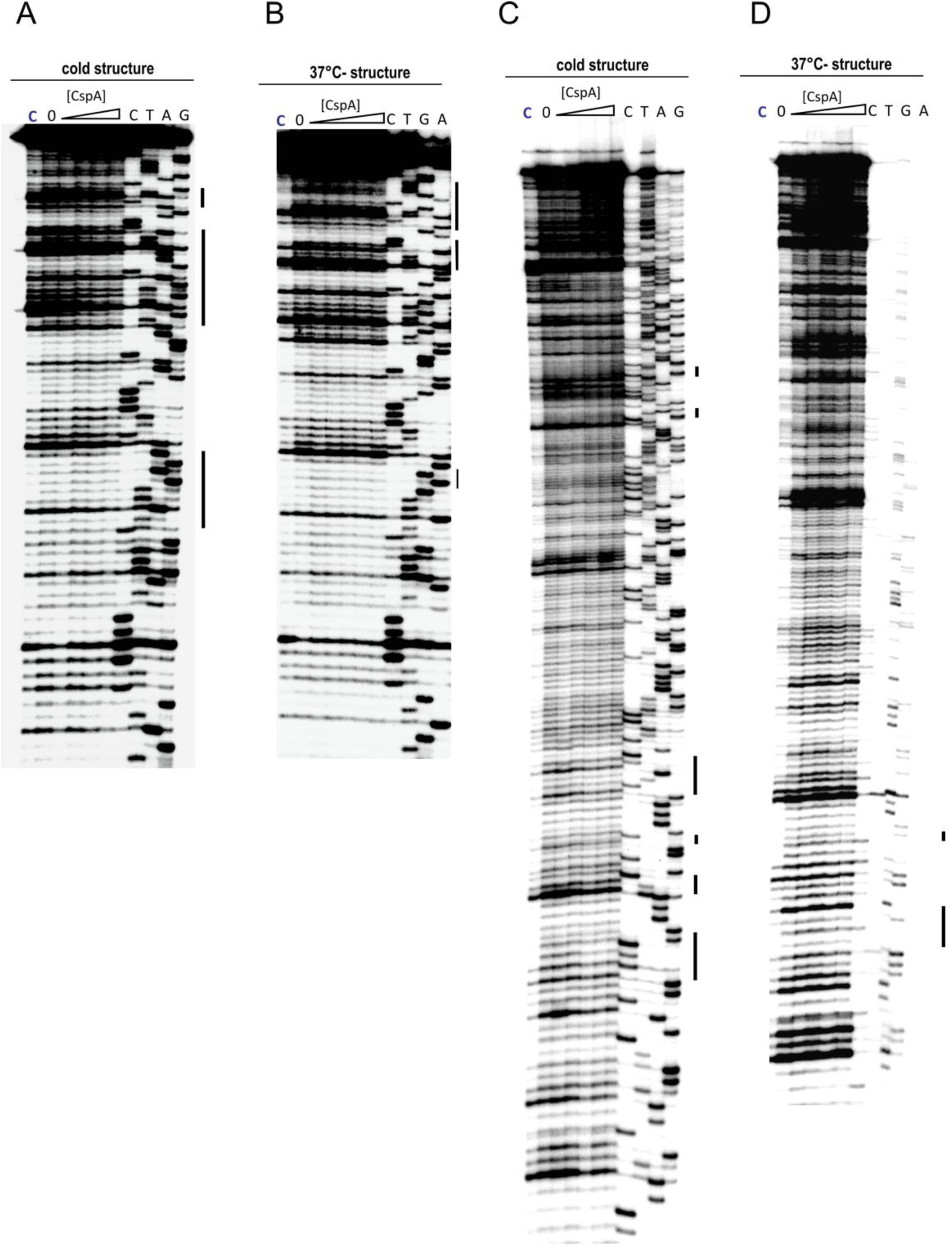
Effect of CspA on the *in situ* accessibility of *cspA* RNA to hydroxyl radical cleavage. Primer extension analysis of the cleavage sites generated by hydroxyl radicals on *cspA* mRNA folded in the conformations indicated at the top of the gels, in the presence of increasing concentrations of CspA (0, 15, 30, 60, 120 μM; increasing concentrations are indicated with a triangle). Lanes C: uncleaved mRNA. Lanes C, T, A, and G correspond to sequencing reactions. Primer extension analysis was performed using primer csp1 (panels A and B), which annealed to the 5′ UTR, and primer csp3 (panels C and D), which annealed to the 3′ UTR. Bases whose accessibility to the cleavage is affected by CspA are highlighted by black bars on the right side of the gels.

**FIGURE 8 supplement 1.**
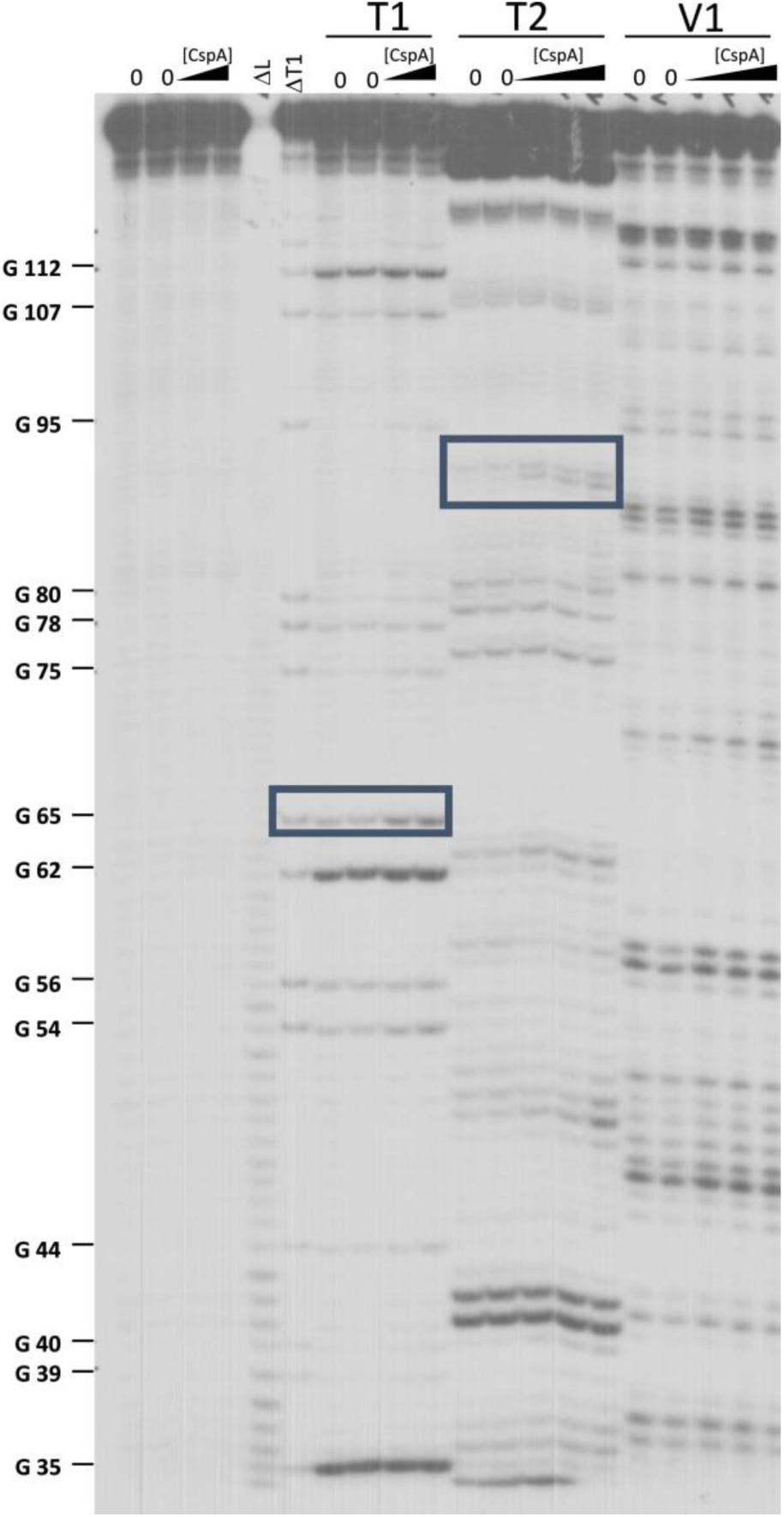
Long electrophoretic migration of the fragments generated by RNase T1 (T1), RNase T2 (T2) or RNase V1 (V1) digestion of 5′-end [^32^P]-labelled 137*cspA* RNA. The experiment was carried out as indicated in Fig. 8.

**FIGURE 10 supplement 1.**
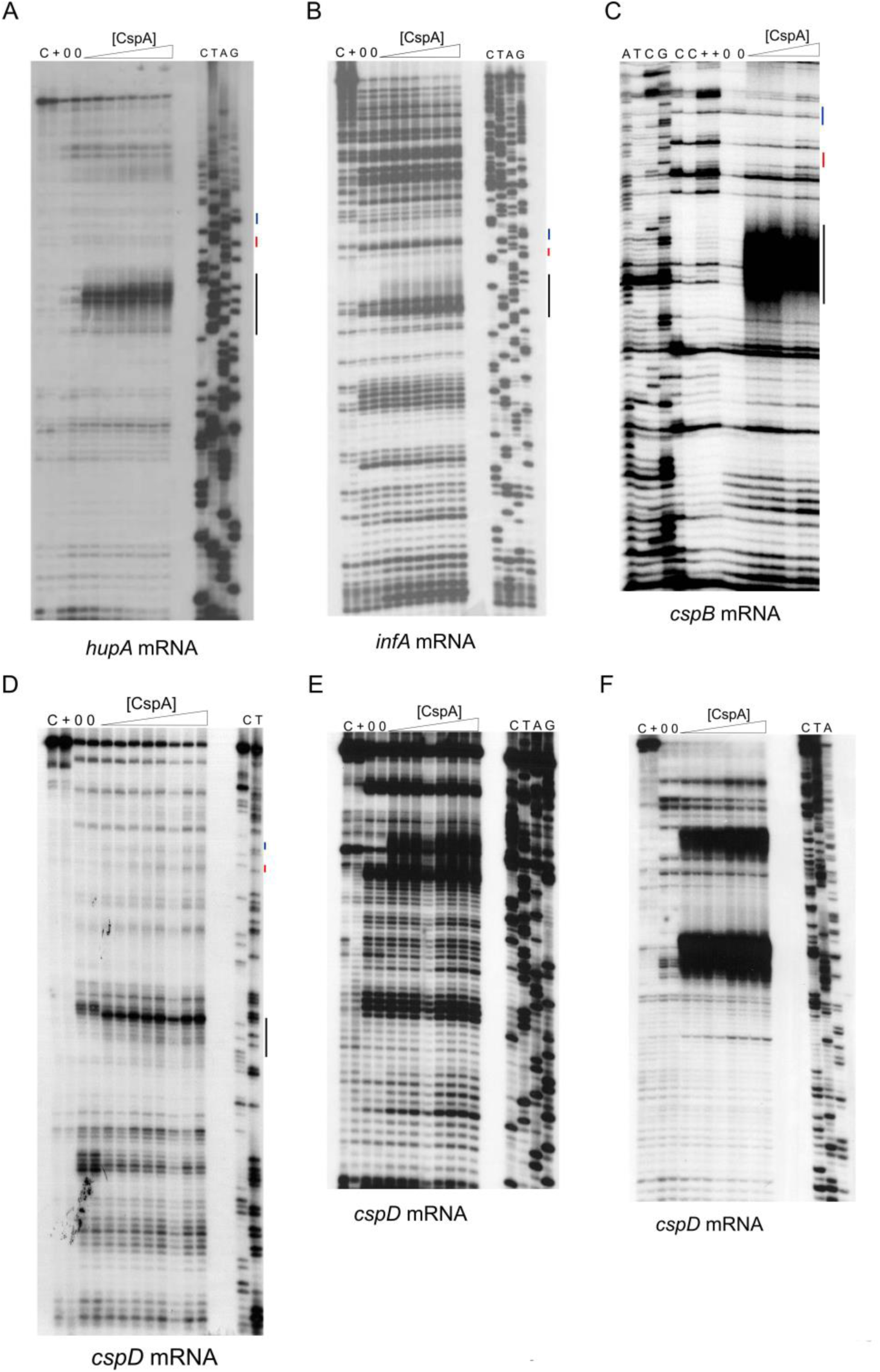
Localization of CspA binding sites on *P1infA, cspB* and *hupA* and *cspD* mRNAs by UV-induced crosslinking at 15°C. The experiment was performed in the presence of 0, 28, 57, and 120 µM of CspA. Lanes C: mRNA alone, no cross-links; lanes +: mRNA+120 µM of CspA, no cross-links. Increasing concentrations are indicated with a triangle. Lanes C, T, A, and G correspond to sequencing reactions. Primer extension analysis was performed: using primers (A) hupA (5′-CTCTGGAGTCCACTCTGGCTG-3′), (B) P1infA (5′-GTTCCGCGTAGAGTTAGAAAACG-3′), and (C) cspB (5′-GGTAGTAAAGATGTGTTTGTG-3′), which annealed 75, 62, and 70 nts downstream from the initiation codon of the corresponding mRNAs, respectively. Primer extension analysis on *cspD* mRNA was performed using primers: (D) cspD3 (5′-CAGATGGATGGTTACAGAACGC-3′), which annealed in the middle of the coding region, (E) cspD4 (5′-GGAAAAGGGTACTGTTAAGTG-3′) which annealed immediately downstream from the initiation codon, and (F) primer cspD2 (5′-GTCTCATTGTGTACATCCTAAAG-3′), which annealed to the 3′ UTR.

## REFERENCES

Andreev, D., Hauryliuk, V., Terenin, I., Dmitriev, S., Ehrenberg, M., Shatsky, I. (2008). The bacterial toxin RelE induces specific mRNA cleavage in the A site of the eukaryote ribosome. RNA 14, 233–239.

Bae, W., Phadtare, S., Severinov, K., and Inouye, M. (1999). Characterization of Escherichia coli cspE, whose product negatively regulates transcription of cspA, the gene for the major cold shock protein. Molecular microbiology 31, 1429–1441.

Barria, C., Malecki, M., and Arraiano, C.M. (2013). Bacterial adaptation to cold. Microbiology 159, 2437–2443.

Brandi, A., Pietroni, P., Gualerzi, C.O., and Pon, C.L. (1996). Post-transcriptional regulation of CspA expression in *Escherichia coli*. Mol. Microbiol. 19, 231–240.

Brandi, A., Spurio, R., Gualerzi, C.O., and Pon, C.L. (1999). Massive presence of the *Escherichia coli* “major cold-shock protein” CspA under non-stress conditions. EMBO J. 18, 1653–1659.

Brandi, A., Giangrossi, M., Giuliodori, A.M., and Falconi, M. (2016) An Interplay among FIS, H-NS, and Guanosine Tetraphosphate Modulates Transcription of the Escherichia coli cspA Gene under Physiological Growth Conditions. Front. Mol. Biosci. 24, 3–19.

Broeze, R.J., Solomon, C.J., and Pope, D.H. (1978). Effects of low temperature on in vivo and in vitro protein synthesis in Escherichia coli and Pseudomonas fluorescens. J Bacteriol. 134, 861–874.

Cristofari, G., and Darlix, J.L. (2002). The ubiquitous nature of RNA chaperone proteins. Progress in Nucleic Acid Research and Molecular Biology 72, 223–268.

Crooks, G.E., Hon, G., Chandonia, J.M., Brenner, S.E. (2004). WebLogo: A sequence logo generator, Genome Research, 14, 1188–1190.

Del Campo, C., Bartholomäus, A., Fedyunin, I., and Ignatova, Z. (2015). Secondary structure across the bacterial transcriptome reveals versatile roles in mRNA regulation and function. PLoS Genet., 11: e1005613.

Di Pietro, F., Brandi, A, Dzeladini, N., Fabbretti, A., Carzaniga, T., Piersimoni, L., Pon, C.L., and Giuliodori, A.M. (2013) Role of the ribosome-associated protein PY in the cold-shock response of Escherichia coli. Microbiologyopen. 2, 293–307.

Donis-Keller, H., Maxam, A.M., and Gilbert, W. (1977). Mapping adenines, guanines, and pyrimidines in RNA. Nucleic Acids Res. 4, 2527–2538.

Dubois, N., Khoo, K.K, Ghossein, S., Seissler, T., Wolff, P., McKinstry, W.J., Mak, J., Paillart, J.C., Marquet, R. and Bernacchi, S. (2018) The C-terminal p6 domain of the HIV-1 Pr55Gag precursor is required for specific binding to the genomic RNA. RNA Biol. 15, 923–936.

Dümmler, A., Lawrence, A.M., and De Marco, A. (2005). Simplified screening for the detection of soluble fusion constructs expressed in *E. coli* using a modular set of vectors. Microb. Cell Fact. 4, 34.

Duval, M., Marenna, A., Chevalier, C., Marzi, S. (2017). Site-Directed Chemical Probing to map transient RNA/protein interactions. Methods. 117, 48–58.

Ermolenko, D.N., and Makhatadze, G.I. (2002). Bacterial cold-shock proteins. Cell Mol Life Sci 59, 1902–1913.

Etchegaray, J.P., and Inouye, M. (1999). A sequence downstream of the initiation codon is essential for cold shock induction of *cspB* of *Escherichia coli*. J Bacteriol. 181, 5852–5854.

Fabbretti, A., Milon, P., Giuliodori, A.M., Gualerzi, C.O., and Pon, C.L. (2007) Real-time dynamics of ribosome-ligand interaction by time-resolved chemical probing methods. Methods Enzymol. 430, 45–58.

Fabbretti, A., Schedlbauer, A., Brandi, L., Kaminishi, T., Giuliodori, A.M., Garofalo, R., Ochoa-Lizarralde, B., Takemoto, C., Yokoyama, S., Connell, S.R., Gualerzi, C.O., and Fucini, P. (2016). Inhibition of translation initiation complex formation by GE81112 unravels a 16S rRNA structural switch involved in P-site decoding. Proc. Natl. Acad. Sci. USA. 113: E2286–95.

Farewell, A., and Neidhardt, F.C. (1998). Effect of temperature on *in vivo* protein synthetic capacity in *Escherichia coli*. J Bacteriol. 180, 4704–4710.

Fang, L., Jiang, W., Bae, W., and Inouye, M. (1997). Promoter-independent cold-shock induction of cspA and its derepression at 37°C by mRNA stabilization. Mol. Microbiol. 23, 355–364.

Fechter, P., Chevalier, C., Yusupova, G., Yusupov, M., Romby, P., and Marzi, S. (2009). Ribosomal initiation complexes probed by toeprinting and effect of trans-acting translational regulators in bacteria. In Riboswitches, Methods in Molecular Biology, A. Serganov, ed. (Totowa, NJ, USA, HumanaPress), pp 247–64.

Friedman, S.M., and Weinstein, I.B. (1964). Lack of fidelity in the translation of synthetic polyribonucleotides. Proceedings of the National Academy of Sciences USA 52, 988–996.

Giangrossi, M., Giuliodori, A.M., Gualerzi, C.O., and Pon, C.L. (2002). Selective expression of the beta-subunit of nucleoid-associated protein HU during cold shock in *Escherichia coli*. Mol Microbiol., 44, 205–16.

Giangrossi, M., Brandi, A., Giuliodori, A.M., Gualerzi, C.O., and Pon, C.L. (2007). Cold-shock-induced *de novo* transcription and translation of *infA* and role of IF1 during cold adaptation. Mol. Microbiol. 64, 807–21.

Giuliodori, A.M., Brandi, A., Gualerzi, C.O., and Pon, C.L. (2004). Preferential translation of cold-shock mRNAs during cold adaptation. RNA 10, 265–276.

Giuliodori, A.M., Giangrossi, M., Brandi, A., Gualerzi, C.O., and Pon, C.L. (2007). Cold-stress-induced *de novo* expression of *infC* and role of IF3 in cold-shock translational bias. RNA 13, 1355–1365.

Giuliodori, A.M., Di Pietro, F., Marzi, S., Masquida, B., Wagner, R., Romby, P., Gualerzi, C.O., and Pon, C.L. (2010) The cspA mRNA is a thermosensor that modulates translation of the cold-shock protein CspA. Mol Cell. 37, 21–33.

Giuliodori, A. M. (2016). Cold-shock response in Escherichia coli: a model system to study post-transcriptional regulation. In Stress and Environmental Regulation of Gene Expression and Adaptation in Bacteria. Frans J. de Bruijn Ed. (New Jersey, USA: Wiley-Blackwell), pp 859–872

Giuliodori, A.M., Fabbretti, A., and Gualerzi, C. (2019) Cold-Responsive Regions of Paradigm Cold-Shock and Non-Cold-Shock mRNAs Responsible for Cold Shock Translational Bias. Int J Mol Sci. 20. pii: E457.

Goldenberg, D., Azar, I., and Oppenheim, A. B. (1996). Differential mRNA stability of the *cspA* gene in the cold-shock response of Escherichia coli. Mol. Microbiol. 19, 241–248.

Goldenberg, D., Azar, I., Oppenheim, A. B., Brandi, A., Pon, C. L., and Gualerzi, C. O. (1997). Role of *Escherichia coli cspA* promoter sequences and adaptation of translational apparatus in the cold-shock response. Mol. Gen. Genet. 256, 282–290.

Goldstein, J.N., Pollitt, S. and Inouye, M. (1990). Major cold shock protein of *Escherichia coli*. Proc. Nati. Acad. Sci. USA 87, 283–287.

Graumann, P., and Marahiel, M.A. (1996) Some like it cold: response of microorganisms to cold shock. Arch Microbiol, 166, 293–300.

Graumann, P.L., and Marahiel, M.A. (1998). A superfamily of proteins that contain the cold-shock domain. Trends Biochem. Sci. 23, 286–90.

Gualerzi, C.O., Giuliodori, A.M., and Pon, C.L. (2003). Transcriptional and post-transcriptional control of cold-shock genes. J. Mol. Biol. 331, 527–539.

Hartz, D., McPheeters, D.S., Traut R., and Gold L. (1988). Extension inhibition analysis of translation initiation complexes. Methods Enzymol. 164, 419–25.

Jiang, W., Hou, Y., and Inouye, M. (1997). CspA, the major cold-shock protein of *Escherichia coli*, is an RNA chaperone. J. Biol. Chem. 272,196–202.

Jones, P.G., Krah, R., Tafuri, S.R., and Wolffe, A.P. (1992). DNA gyrase, CS7.4, and the cold shock response in *Escherichia coli*. J Bacteriol. 174, 5798–5802.

Jossinet, F., Ludwig, T.E., and Westhof, E. (2010). Assemble: an interactive graphical tool to analyze and build RNA architectures at the 2D and 3D levels. Bioinformatics 26, 2057–2059.

Kapust, R. B., Tözsér, J., Copeland, T.D., and Waugh, D.S. (2002) The P1’ specificity of tobacco etch virus protease. Biochem. Biophys. Res. Commun. 5, 949–955.

Kremer, W., Schuler, B., Harrieder, S., Geyer, M., Gronwald, W., Welker, C., Jaenicke, R., and Kalbitzer, H.R. (2001). Solution NMR structure of the cold-shock protein from the hyperthermophilic bacterium Thermotoga maritima. Eur. J. Biochem. 268, 2527–2539.

Liu, T., Kaplan, A., Alexander, L., Yan, S., Wen, J.D., Lancaster, L., Wickersham, C.E., Fredrick, K., Noller, H., Tinoco, I., et al. (2014). Direct measurement of the mechanical work during translocation by the ribosome. eLife 3, e03406.

La Teana, A., Brandi, A., Falconi, M., Spurio, R., Pon, C. L., and Gualerzi, C. O. (1991). Identification of a cold shock transcriptional enhancer of the *Escherichia coli* gene encoding nucleoid protein H-NS. Proc. Natl Acad. Sci. USA, 88, 10907–10911.

Lopez, M.M., and Makhatadze, G.I. (2000). Major cold shock proteins, CspA from *Escherichia coli* and CspB from *Bacillus subtilis*, interact differently with single-stranded DNA templates. Biochim Biophys Acta, 1479, 196–202.

Mayer, O., Rajkowitsch, L., Lorenz, C., Konrat, R., and Schroeder, R. (2007). RNA chaperone activity and RNA-binding properties of the E. coli protein StpA. Nucleic Acids Res. 35, 1257–1269.

Mitta, M., Fang, L., and Inouye, M. (1997). Deletion analysis of *cspA* of *Escherichia coli*: requirement of the AT-rich UP element for *cspA* transcription and the downstream box in the coding region for its cold shock induction. Mol Microbiol 26, 321–335.

Mueller, U., Perl, D., Schmid, F.X., and Heinemann, U. (2000). Thermal stability and atomic-resolution crystal structure of the Bacillus caldolyticus cold shock protein. J. Mol. Biol. 297, 975–988.

Newkirk, K., Feng, W., Jiang, W., Tejero, R., Emerson, S.D., Inouye, M., and Montelione, G.T. (1994). Solution NMR structure of the major cold shock protein (CspA) from Escherichia coli: identification of a binding epitope for DNA. Proc. Natl Acad. Sci. USA 91, 5114–5118.

Neubauer, C., Gao, Y-G., Andersen, K.R., Dunham, C.M., Kelley, A.C., Hentschel, J., Gerdes, K., Ramakrishnan, V., Brodersen, D.E. (2009). The Structural Basis for mRNA Recognition and Cleavage by the Ribosome-Dependent Endonuclease RelE. Cell. 139, 1084–1095.

Pedersen, K., Zavialov, A.V., Pavlov, M.Y., Elf, J., Gerdes, K., Ehrenberg, M. (2003). The bacterial toxin RelE displays codon-specific cleavage of mRNAs in the ribosomal A site. Cell. 112, 131–140.

Phadtare, S., and Inouye, M. (1999). Sequence-selective interactions with RNA by CspB, CspC and CspE, members of the CspA family of *Escherichia coli*. Mol Microbiol 33, 1004– 1014.

Phadtare, S., and Inouye, M. (2004). Genome-wide transcriptional analysis of the cold shock response in wild-type and cold-sensitive, quadruple-csp-deletion strains of *Escherichia coli*. J Bacteriol. 186, 7007–7014.

Phadtare, S. (2004). Recent developments in bacterial cold-shock response. Curr. Issues Mol. Biol. 6, 125–36.

Phadtare, S., and Severinov, K. (2009) Comparative analysis of changes in gene expression due to RNA melting activities of translation initiation factor IF1 and a cold shock protein of the CspA family. Genes Cells. 14, 1227–39.

Qu, X., Wen, J.D., Lancaster, L., Noller, H.F., Bustamante, C., and Tinoco, I., Jr. (2011). The ribosome uses two active mechanisms to unwind messenger RNA during translation. Nature 475, 118–121.

Rajkowitsch, L., and Schroeder R. (2007). Dissecting RNA chaperone activity. RNA 13, 2053–60.

Rennella, E., Sara, T., Juen, M., Wunderlich, C., Imbert, L., Solyom, Z., Favier, A., Ayala, I., Weinhaupl, K., Schanda, P., et al. (2017). RNA binding and chaperone activity of the E. coli cold-shock protein CspA. Nucleic Acids Res. 45, 4255–4268.

Sachs, R., Max, K.E., Heinemann, U., Balbach, J. (2012). RNA single strands bind to a conserved surface of the major cold shock protein in crystals and solution. RNA, 18, 65–76.

Serganov, A., Rak, A., Garber, M., Reinbolt, J., Ehresmann, B., Ehresmann, C., Grunberg-Manago, M., and Portier, C. (1997). Ribosomal protein S15 from *Thermus thermophilus*--cloning, sequencing, overexpression of the gene and RNA-binding properties of the protein. Eur. J. Biochem. 246, 291–300.

Shimizu, Y., Inoue, A., Tomari, Y., Suzuki, T., Yokogawa, T., Nishikawa, K., and Ueda, T. (2001). Cell-free translation reconstituted with purified components. Nat. Biotechnol. 19, 751–755.

Schindelin, H., Jiang, W., Inouye, M., and Heinemann, U. (1994). Crystal structure of CspA, the major cold shock protein of *Escherichia coli*. Proc. Natl Acad. Sci. USA, 91, 5119–5123.

Schnuchel, A., Wiltscheck, R., Czisch, M., Herrler, M., Willimsky, G., Graumann, P., Marahiel, M.A., and Holak, T.A. (1993). Structure in solution of the major cold-shock protein from Bacillus subtilis. Nature 364, 169–171.

Weber, M.H., and Marahiel, M.A. (2003) Bacterial cold shock responses. Sci. Prog., 86, 9– 75.

Withman, B., Gunasekera, T.S., Beesetty, P., Agans, R. and Paliy, O. (2013) Transcriptional responses of uropathogenic *Escherichia coli* to increased environmental osmolality caused by salt or urea. Infect Immun., 81, 80–89.

Wolffe, A.P., Tafuri, S., Ranjan, M., and Familari, M. (1992). The Y-box factors: a family of nucleic acid binding proteins conserved from *Escherichia coli* to man. New Biol. 4, 290–298.

Yamanaka, K., and Inouye, M. (1997). Growth-phase-dependent expression of *cspD*, encoding a member of the CspA family in *Escherichia coli*. J Bacteriol., 179, 5126–5130.

Yamanaka, K., Fang, L., and Inouye, M. (1998) The CspA family in *Escherichia coli*i: multiple gene duplication for stress adaptation. Mol. Microbiol., 27, 247–255.

Yamanaka, K., Mitta, M., and Inouye, M. (1999). Mutation analysis of the 5’-untranslated region of the cold shock *cspA* mRNA of Escherichia coli. J Bacteriol. 181, 6284–6291.

Xia, B., Ke, H., and Inouye, M. (2001). Acquirement of cold-sensitivity by quadruple deletion of the *cspA* family and its suppression by PNPase S1 domain in *Escherichia coli*. Mol Microbiol, 40, 179– 188.

Zhang, Y., Burkhardt, D.H., Rouskin, S., Li, G.W., Weissman, J.S., and Gross C.A. (2018). A Stress Response that Monitors and Regulates mRNA Structure Is Central to Cold Shock Adaptation. Mol Cell, 70, 274–286

Zuker, M. (2003). Mfold web server for nucleic acid folding and hybridization prediction. Nucleic Acids Res. 31, 3406–15.

